# Muscle-directed mechanosensory feedback activates egg-laying circuit activity and behavior in *C. elegans*

**DOI:** 10.1101/2022.07.19.500701

**Authors:** Emmanuel Medrano, Kevin M. Collins

## Abstract

Mechanosensory feedback of internal reproductive state drives decisions about when and where to reproduce.^1^ For instance, stretch in the *Drosophila* reproductive tract produced by artificial distention or from accumulated eggs regulates the attraction to acetic acid to ensure optimal oviposition.^2^ How such mechanosensory feedback modulates neural circuits to coordinate reproductive behaviors is incompletely understood. We previously identified a stretch-dependent homeostat that regulates egg laying in *Caenorhabditis elegans*. Sterilized animals lacking eggs show reduced Ca^2+^ transient activity in the presynaptic HSN command motoneurons that drive egg-laying behavior while animals forced to accumulate extra eggs show dramatically increased circuit activity that restores egg laying.^3^ Interestingly, genetic ablation or electrical silencing of the HSNs delays, but does not abolish, the onset of egg laying^3–5^ with animals recovering vulval muscle Ca^2+^ transient activity upon egg accumulation.^6^ Using an acute gonad microinjection technique to mimic changes in pressure and stretch resulting from germline activity and egg accumulation, we find that injection rapidly stimulates Ca^2+^ activity in both neurons and muscles of the egg-laying circuit. Injection-induced vulval muscle Ca^2+^ activity requires L-type Ca^2+^ channels but is independent of presynaptic input. Conversely, injection-induced neural activity is disrupted in mutants lacking the vulval muscles, suggesting ‘bottom-up’ feedback from muscles to neurons. Direct mechanical prodding activates the vulval muscles, suggesting they are the proximal targets of the stretch-dependent stimulus. Our results show that egg-laying behavior in *C. elegans* is regulated by a stretch-dependent homeostat that scales postsynaptic muscle responses with egg accumulation in the uterus.

## Results and Discussion

While microinjection into the gonad of the *C. elegans* is a common technique for generating transgenic strains,^7,8^ we noticed injection often led to vulval opening and egg release. While we feel this phenomenon is widely known among *C. elegans* researchers, it is not well-described in the literature. To determine if microinjection promotes egg laying, we inserted a needle into the syncytial gonad of wild-type animals and performed a brief (3 s) injection delivering a median volume of ~250 pL of injection buffer (see STAR methods) while monitoring egg release (**Figure 1A**). Microinjection drove egg release in nearly half (~46 ± 9%) of animals (**Figure 1B**) but not in *unc-54(e190)* muscle-specific myosin mutants which cannot contract the vulval muscles (**Figure 1B**).^9^

**Figure 1:**
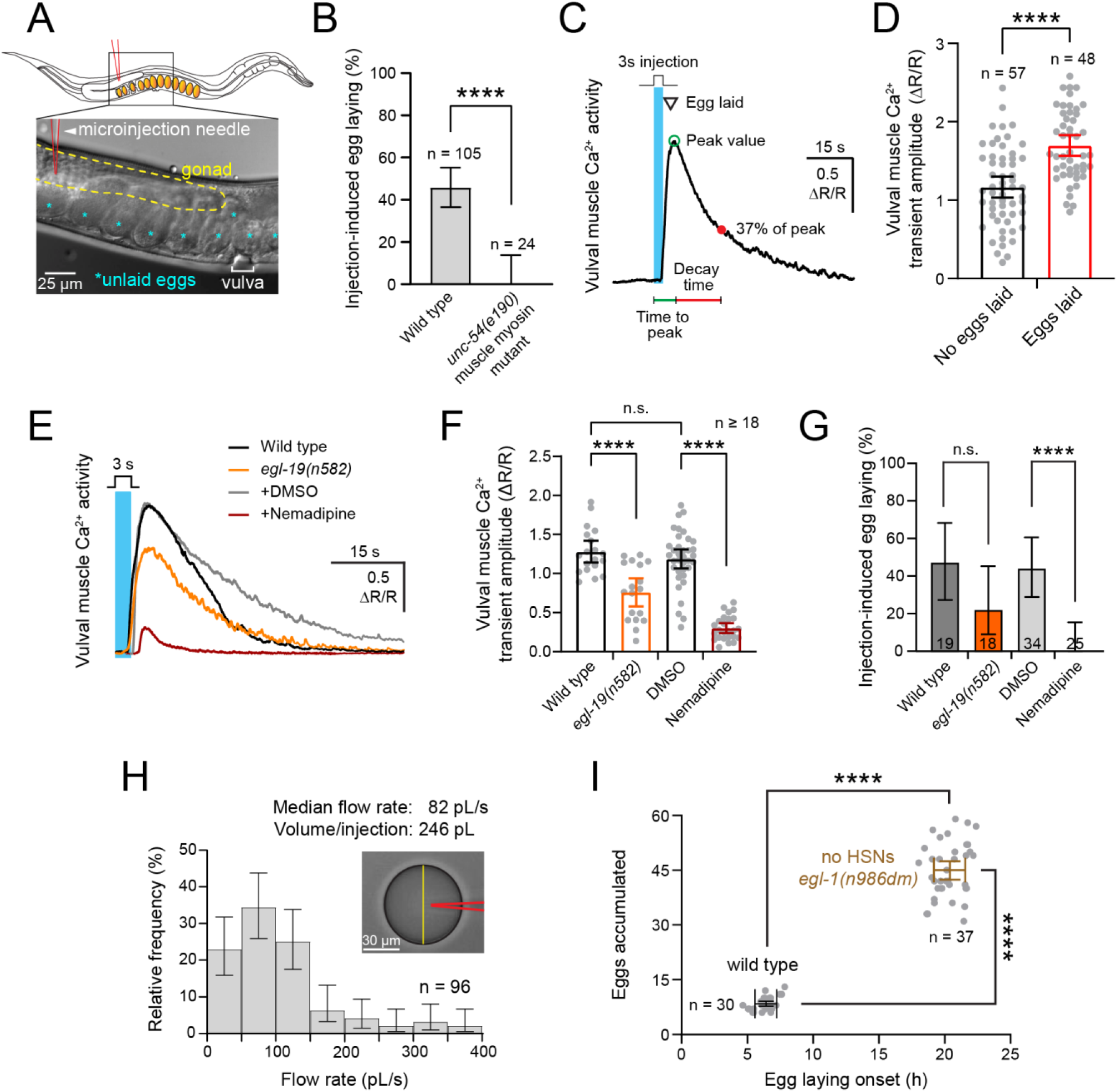
Acute microinjection triggers vulval muscle Ca^2+^ activity and egg-laying behavior. **(A)** Schematic of *C. elegans* and micrograph demonstrating the injection technique. A microinjection needle (arrowhead; red outline) is inserted into the gonad (yellow outline) of an immobilized *C. elegans* worm (top). A 3 s pulse is applied to inject a standard microinjection buffer into the gonad syncytium. Unlaid eggs (cyan asterisks) are then monitored for injection-induced egg release through the vulva (white bracket). **(B)** Bar graphs showing the percent of wild-type or *unc-54(e190)* muscle myosin mutants animals laying eggs in response to injection. Error bars represent ± 95% confidence interval for proportions; asterisks indicate p-value < 0.0001 (Fisher’s exact test, n indicates number of animals injected per genotype). **(C)** Representative vulval muscle Ca^2+^ trace, determined from GCaMP5/mCherry fluorescence ratio, following a 3 s injection pulse (vertical cyan bar). Key features observed from a typical injection response are egg release (inverted triangle), peak amplitude (ΔR/R; open green circle), time to peak (relative to start of injection; green horizontal line), and decay time (s; time from peak amplitude to 37% peak value; red horizontal line). **(D)** Bar graphs comparing mean vulval muscle Ca^2+^ transient amplitudes (± 95% confidence interval) after injections that did not (black) and those that did (red) result in egg release. Points in scatterplot show responses in individual animals. Asterisks indicate p-value < 0.0001 (unpaired t-test). **(E)** Representative injection-induced vulval muscle Ca^2+^ responses for wild-type (black) or *egl-19(n582)* voltage-gated Ca^2+^ channel loss-of-function mutant animals (orange), worms exposed to 0.1% DMSO (grey), and worms exposed to 25 µM nemadipine (burgundy). Vertical cyan bar represents 3 s injection pulse. **(F)** Bar graphs with scatterplot of mean vulval muscle Ca^2+^ transient amplitude in wild-type (black), *egl-19(lf)* (orange), 0.1% DMSO-treated worms (grey), and nemadipine-treated animals (burgundy) after injection. Asterisks indicates p-value < 0.0001 and n.s. indicates p-value > 0.05 (one-way ANOVA with Bonferroni’s correction for multiple comparisons; n ≥ 18 animals). **(G)** Bar graphs showing percent of injections that resulted in an egg-laying event (mean ± 95% confidence intervals for the proportion). Numbers inside bar graph indicate sample sizes; n ≥ 18 animals per condition. Asterisks indicate p-value < 0.0001; n.s. p-value > 0.05 (Pairwise Fisher’s exact test). **(H)** Distribution of estimated injection flow rates obtained by measuring the change in diameter over time of injection buffer drops (yellow vertical line in inserted micrograph) into halocarbon oil immediately following needle withdrawal after worm injection. Error bars represent ± 95% confidence interval for the mean proportion across 96 injections. **(I)** Scatterplot of time to egg-laying onset and number of eggs accumulated at egg-laying onset in wild-type (black) and *egl-1(n986dm)* HSN-deficient mutant animals (brown). Asterisks indicate p-value < 0.0001 (unpaired t-tests for differences in egg-laying onset and in egg accumulation).

As muscle contraction requires Ca^2+^ activity,^10–12^ we performed ratiometric GCaMP5 recordings of the vulval muscles.^6^ Gonad injections triggered a strong and rapid induction of vulval muscle Ca^2+^ activity which peaked on average 5.6 s after the onset of injection (**Figure 1C; Video S1**) with subsequent injections showing weaker induction of Ca^2+^ (**Figure S1A**). Direct contact with the injected buffer did not seem to be required for a robust muscle response, as Ca^2+^ activity was induced immediately upon injection while the buffer reached the vulva a few seconds later (**Figure S2A**). As seen in behaving animals,^6,10^ stronger vulval muscle Ca^2+^ transients drove egg-laying events (**Figure 1D**) in wild-type but not *unc-54(e190)* mutant animals (**Figure S1B**). The number of eggs in the uterus prior to injection correlated significantly with the resulting Ca^2+^ peak amplitude elicited by injections (**Figure S1C**), with higher Ca^2+^ transient amplitudes driving egg laying (**Figure 1D**), consistent with a model where stretch caused by embryo accumulation enhances vulval muscle excitability and the likelihood of egg laying.

Ca^2+^ entry into the vulval muscles is mediated by EGL-19 L-type voltage-gated Ca^2+^ channels.^11,13–15^ To test if the injection response requires L-type Ca^2+^ channels, we recorded vulval muscle responses after channel function was blocked genetically or pharmacologically (**Figure 1E**). *egl-19(n582)* loss-of-function mutants, or wild-type animals treated for ~3 hours with 25 µM nemadipine, a specific blocker of L-type channels,^15^ showed significantly reduced vulval muscle Ca^2+^ responses following injection (**Figure 1E – F**; **Video S2**). Injections also failed to elicit egg laying in nemadipine-exposed animals when compared to DMSO-treated worms (**Figure 1G**). As such, injection-induced vulval muscle Ca^2+^ and egg-laying requires L-type Ca^2+^ channels. Together, these results show that microinjection does not simply force eggs out through the vulva. Instead, microinjection induces an active response, triggering a rapid onset of vulval muscle Ca^2+^ activity, muscle contractility, and egg release.

To determine if the magnitude of stretch affected the vulval muscle Ca^2+^ response, we estimated injection flow rates across multiple animals to determine total volume delivered per injection. Injection flow rates ranged between ~10 and ~366 pL/s, corresponding to a total volume delivered ranging between 30 pL – 1 nL, with a median of 246 pL for each 3 s injection (**Figure 1H**). As each *C. elegans* embryo is ~25 pL, each injection delivered an equivalent stretch of 1.2 – 40 eggs (median ~10 egg-volumes per injection). To determine how this range of injection volumes compares to the number of eggs required to activate the stretch-dependent homeostat, we measured egg accumulation at the onset of egg laying in wild-type or *egl-1(dm)* HSN-deficient mutants whose vulval muscle Ca^2+^ activity depends upon feedback of egg accumulation.^3,6^ Wild-type animals begin laying eggs ~6 hours after reaching adulthood, having accumulated ~8 eggs in the uterus (**Figure 1I**). Loss of HSNs delays egg laying to ~20 hours after ~45 eggs have accumulated (**Figure 1I**). We interpret this difference in ~37 eggs as providing a threshold estimate of stretch and/or volume increase (~925 pL) required to activate the homeostat in the absence of HSN input. As such, the volumes injected here are quantitatively similar to the volume changes that accompany egg production and induce egg laying *in vivo*.

We explored additional parameters that affected the injection response including ionic composition of injection buffer, volume injected, injection flow rate, site of injection into the gonad, and age and fertility of injected animals (**Figure S1D – H; Figure S2; see STAR Methods**). Within typical injections performed between 30 – 40 psi, there was no significant correlation between flow rate and vulval muscle Ca^2+^ (**Figure S1D**). However, at lower flow rates, we found that ‘fast’ injections’ drove stronger vulval muscle Ca^2+^ transients compared to ‘slow’ injections of similar volume (**Figure S2C – E; See STAR Methods**). Injections caused a 10% average increase in relative body size that correlated with the vulval muscle Ca^2+^ response (**Figure S1I**). Injections also caused movement of the vulval muscles within the body **(Figure S1J**). Thus, injection causes an immediate change in animal size along with vulval muscle position and Ca^2+^ activity, possibly reflecting the natural processes by which egg accumulation and/or positioning stimulate the egg-laying homeostat.^3–5^ Since genetic or chemical sterilization of adult animals prevents egg-laying circuit activity,^3^ we tested whether microinjection into these animals would mimic feedback of egg accumulation and rescue Ca^2+^ activity. Injections into FUDR-sterilized,^16^ *fog-2* sperm-deficient^17^ adults, or juvenile fourth-larval stage animals triggered strong vulval muscle Ca^2+^ transients that were comparable to wild-type and fertile adults (**Figure S3**). These results suggest stretch-dependent feedback–not animal age, fertility, or even the eggs themselves–is sufficient to activate vulval muscle Ca^2+^ activity and contractility for egg release.

Egg laying is driven by a rhythmic and sequential pattern of Ca^2+^ activity in the neurons and muscles of the egg-laying circuit. The serotonergic HSN command neurons go first, with Ca^2+^ transients peaking ~2 s before each egg-laying event during which Ca^2+^ transients are observed in the VC motoneurons and the vulval muscles.^6^ The tyraminergic uv1 neuroendocrine cells show Ca^2+^ activity last, typically ~2 s later, in response to vulval opening and egg release (**Figure 2A**). To determine if injection drives a similar pattern of Ca^2+^ activity, we injected animals expressing GCaMP5 in either the HSNs, VCs, or uv1 cells. All cells of the circuit showed a robust induction of Ca^2+^ activity following injection, but their order and kinetics of onset differed from that seen in behaving animals. The vulval muscles and the proximally located uv1 cells showed the most rapid responses (**Video S3**), reaching half-maximal Ca^2+^ levels at 2.4 ± 1.2 and 3.4 ± 3.3 s (mean ± SD) after injection onset, respectively (**Figure 2B and 2C**). The VCs followed, showing half-maximal Ca^2+^ activity at 4.0 ± 1.5 s (**Figure 2D**; **Video S4**). Surprisingly, the HSNs showed the slowest response at 6.4 ± 4.0 s (**Figure 2E and 2F**; **Video S5**). Injection-induced Ca^2+^ responses typically lasted longer and decayed more slowly compared to those seen in behaving animals which may result from the absence of rhythmic phasing of Ca^2+^ activity that accompanies locomotion.^6,10,11^ The robust and rapid onset of vulval muscle Ca^2+^ activity upon microinjection, and subsequent induction of uv1, VC, and HSN Ca^2+^ activity (**Figure 2F**), suggests the vulval muscles are the initial responders.

**Figure 2:**
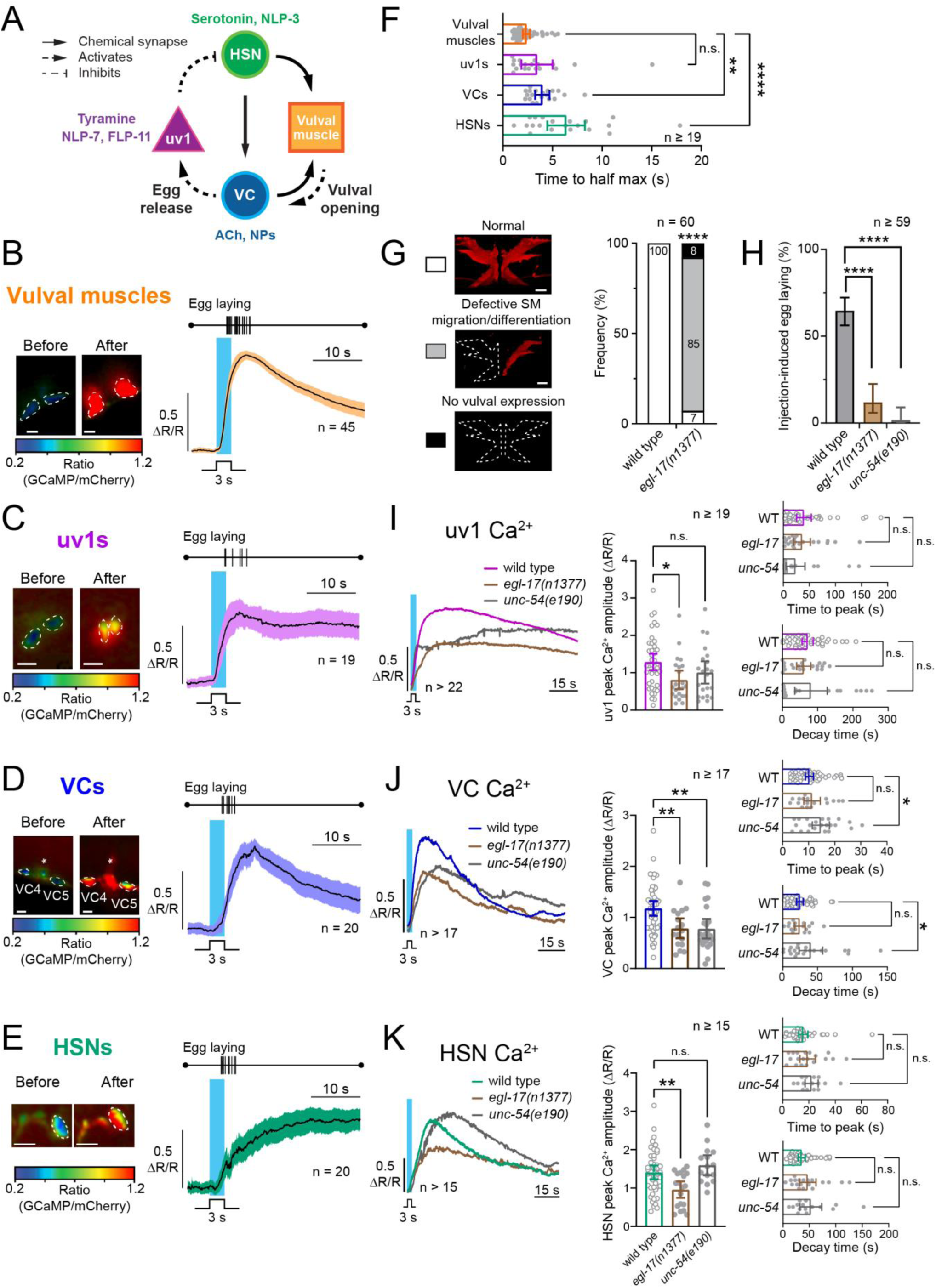
The vulval muscles are necessary for the sequential Ca^2+^ activity of the egg-laying circuit induced by microinjections. **(A)** Connectivity of the *C. elegans* egg-laying circuit^6^ The HSNs promote vulval muscle activity via serotonin and NLP-3 signaling^5,55,56^ which in turn leads to muscle contractions that promote VC activity^19^. The VCs also promote muscle contractions via acetylcholine and neuropeptide signaling^5,6^. Thereafter VC Ca^2+^ activity promotes successful egg laying which leads to the mechanical stimulation of the uv1 cells (dotted arrow) that act to inhibit the HSNs and terminate egg-laying via signaling through tyramine, NLP-7, and FLP-11^4,12,22,25^. **(B – E)** Normalized injection-induced Ca^2+^ response of the vulval muscles (B), uv1 neuroendocrine cells (C), VC neurons (D), and HSN neurons (E). Vertical cyan line shows 3 s injection period. Micrographs show Ca^2+^ activity in cell bodies (outlined in white) before and after injection; rainbow scale indicates GCaMP5/mCherry ratio in responding cells; blue indicates low Ca^2+^ and red indicates elevated Ca^2+^. White scale bar = 10 µm. Black solid line is a trace of the average Ca^2+^ response; colored bands indicate the 95% confidence interval (n ≥ 19 animals per trace). Individual cell Ca^2+^ responses were normalized by minimal and maximal response following injection. **(F)** Bar graphs with scatter plots showing time of half-max Ca^2+^ responses. **** indicates p-value < 0.0001, ** indicates p-value = 0.0013, and n.s. indicates p-value > 0.05 (Kruskal-Wallis test with Dunn’s correction for multiple comparisons). Vertical tick marks show timing of egg-laying events pooled from all injected animals. **(G)** Micrographs demonstrating the three categories of vulval muscle mCherry expression found in wild-type or *egl-17(n1377)* mutant animals with defective sex myoblast migration and/or differentiation (left). White scale bar = 10 µm. Relative percentage of wild type or *egl-17* animals showing each type of expression pattern (right). Number within bar graph represents relative percentage within category, while **** represents p-value < 0.0001 (Fisher’s exact test comparing normal and abnormal vulval muscle development). **(H)** Bar graphs showing percent of wild type, *egl-17*, or *unc-54* mutant animals that laid eggs following microinjection (error bars represent ± 95% C.I.). **(I – K)** Average injection-induced Ca^2+^ responses from the uv1s (I), VCs (J), and HSNs (K) in wild-type (open circles), *egl-17(n1377)* (brown), or *unc-54(e190)* (grey) mutant animals. Bar graphs with scatterplots show average peak amplitudes (middle), time to peak (top right), and decay times (bottom right) for the uv1, VC, or HSN cells. Error bars represent ± 95% confidence intervals, n.s. = not significant (p-value > 0.05), * indicates p-value < 0.05, ** indicates p-value < 0.01 (ANOVA with Bonferroni’s correction for multiple comparisons).

To test whether the vulval muscles transmit the injection response to the rest of the circuit, we injected into *egl-17(n1377)* mutants that lack vulval muscle development due to improper sex myoblast migration.^18^ 85% of *egl-17(n1377)* mutant animals lacked the anterior vulval muscles and 8% lacked both anterior and posterior muscles (**Figure 2G**), preventing egg release similarly to *unc-54* muscle myosin mutants (**Figure 2H**). *egl-17* mutants also showed a significant reduction in average Ca^2+^ peak amplitudes in uv1s, VCs, and HSNs following injection (**Figure 2I – K**), suggesting the vulval muscles are required for their activity. *unc-54* mutants showed a similar reduction only in VC Ca^2+^ activity along with delayed rise and decay kinetics (**Figure 2J**), consistent with previous results showing that VC Ca^2+^ activity is induced by vulval muscle contractility following optogenetic stimulation.^19^ As injection drives robust vulval muscle Ca^2+^ activity in *unc-54* mutants (**Figure S1B**), release of retrograde factors may facilitate the activation of HSNs and uv1s.^3^ Together, these results show the vulval muscles are required, at least in part, for activation of other cells in the egg-laying circuit, suggesting the vulval muscles initiate the injection response.

To test the hypothesis that the vulval muscles respond cell-autonomously, we removed synaptic and peptidergic input into the vulval muscles and tested their Ca^2+^ response to microinjection. Specifically, we tested *egl-1(dm)* mutant animals lacking HSNs and in which we blocked synaptic transmission from the VCs through expression of Tetanus Toxin (**Figure 3A**).^19^ The vulval muscles of these animals showed strong injection-induced Ca^2+^ activity (**Figure 3B**), with no significant differences in Ca^2+^ responses or egg laying compared to wild-type animals (**Figure 3C – F**). These results indicate the main synaptic inputs into the vulval muscles are not required for their injection response, consistent with previous work showing that worms lacking HSNs and/or VCs still enter egg-laying active states^5^ with robust vulval muscle Ca^2+^ activity.^6,11,12,19^ To account for the possibility that other cells, including those outside the egg-laying circuit, respond to acute injection and activate the vulval muscles, we injected *unc-13(s69)* mutants lacking synaptic vesicle transmission^20^ or *unc-31(e928)* mutants with defective dense core vesicle release.^21^ Both mutants showed normal vulval muscle Ca^2+^ responses after microinjection (**Figure 3B – E**) and were just as likely to lay eggs (**Figure 3F**). Because injection induces vulval muscle and uv1 Ca^2+^ activity with similar kinetics (**Figure 2F**), we wanted to test whether peptidergic or tyramine signals from uv1 are required (**Figure 3A**).^22^ Injection into *egl-3(ok979)* or *egl-21(n476)* mutants in which proneuropeptide synthesis is disrupted,^23,24^ or into *tdc-1* mutants lacking tyramine,^25^ showed normal induction of vulval muscle Ca^2+^ activity and egg laying (**Figure 3B – F**). Together, these results show that the vulval muscles respond to injection independent of synaptic or peptidergic signals, consistent with these mutants laying eggs at least under some laboratory growth conditions,^26,27^ suggesting feedback of germline activity and/or egg accumulation activates the muscles.

**Figure 3:**
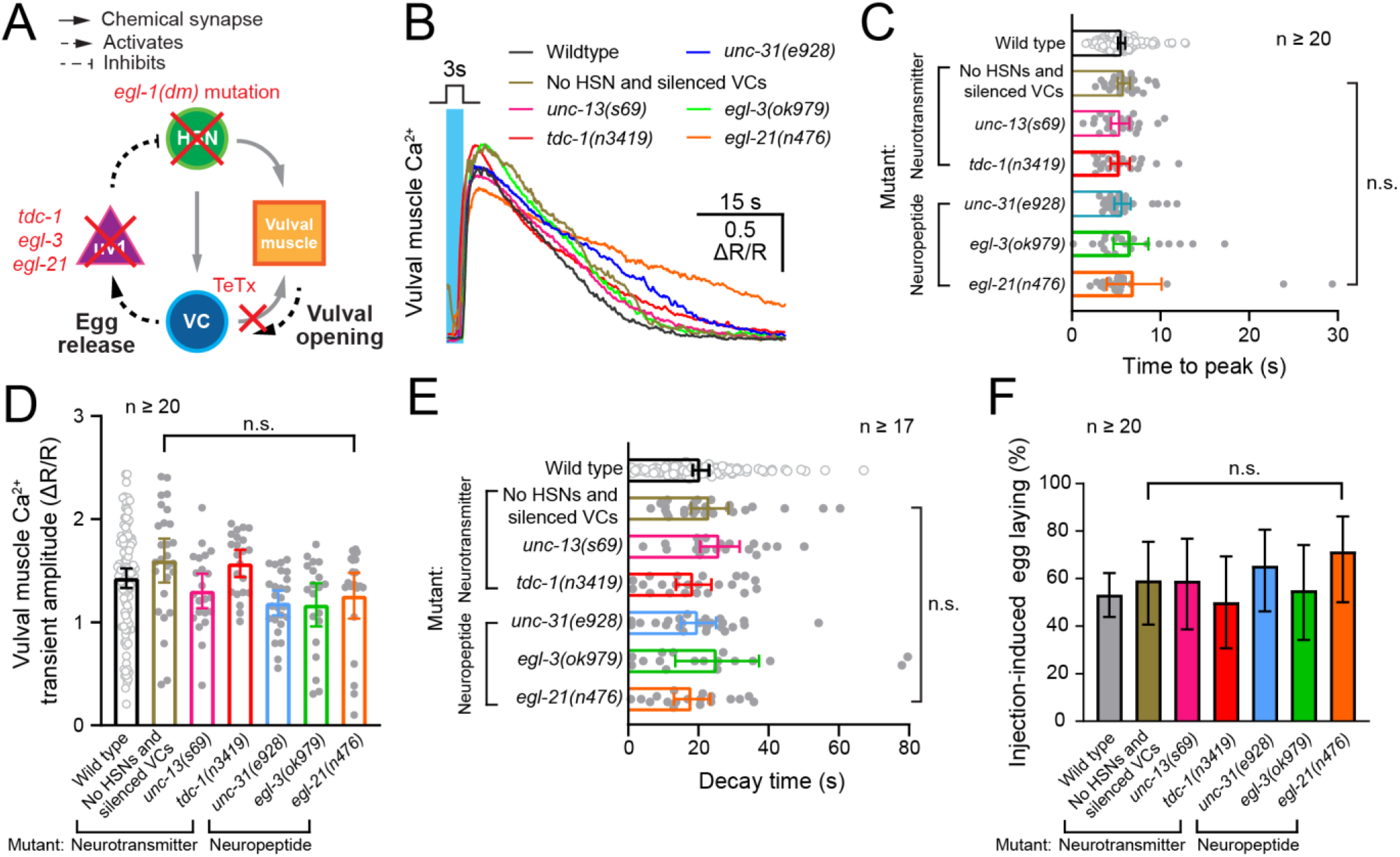
The vulval muscle Ca^2+^ and egg-laying response to microinjection does not require synaptic or peptidergic input. **(A)** Diagram highlighting the synaptic connections between cells of the *C. elegans* egg-laying circuit. Red “x” symbol on top of the HSNs represents the genetic removal of any input from the HSNs cells through use of *egl-1(n986dm)* mutants in which the HSN neurons undergo programmed cell death. Tetanus Toxin (TeTx) was also transgenically expressed in the VC neurons to block their neurotransmitter and neuropeptide release. Grayed-out arrows represent loss of synaptic communication in mutants with no HSNs and silenced VCs. Red “x” on uv1s represents the silencing of uv1s via neurotransmitter or neuropeptide mutants using *tdc-1(n3419), egl-3(ok979), or egl-21(n476)* animals. **(B)** Representative injection-induced vulval muscle Ca^2+^ responses of wild-type (black), HSN-deficient *egl-1(dm)* mutants co-expressing Tetanus Toxin in the VCs (brown), *unc-13(s69)* mutants with defective synaptic transmission, (magenta), *tdc-1(n3419)* mutants with defective tyramine synthesis (red), *unc-31(e928)* mutant animals with defective peptidergic transmission (blue) or *egl-3(ok979)* and *egl-21(n476)* mutants defective in peptide synthesis (green and orange, respectively). Vertical cyan bar represents the 3 s gonad injection. **(C – E)** Bar graphs with scatter plot showing mean time to vulval muscle Ca^2+^ transient peak (C), mean Ca^2+^ transient peak amplitude (D), and mean Ca^2+^ transient decay time (E). Open circles represent wild type. **(F)** Bar graphs showing percent of injections that led to an egg-laying event (mean ± 95% confidence intervals for the proportion); n.s. indicates p-value > 0.05 (Chi-square test).

If the vulval muscles are sensitive to stretch, they may respond to other forms of mechanical stimulation. We used a glass capillary probe to prod the vulval muscles while recording changes in muscle Ca^2+^ activity (**Figure 4A**). Mechanical stimulation of the vulval muscles triggered a strong and immediate Ca^2+^ activity in the prodded muscle (**Video S6**). We subjected animals to a train of prodding events (up to 35 µm; **Figure 4B**), finding a significant correlation between vulval muscle displacement and maximal Ca^2+^. We then applied 1 s, 30 µm prodding stimulations to the vulval muscles and compared Ca^2+^ responses in anterior/posterior pairs where only one set of muscles received the stimulus. We observed stronger Ca^2+^ transients in the prodded muscles (**Figure 4C – D**), with a half-maximal Ca^2+^ response time of 0.7 ± 0.4 s (mean ± SD). This response was faster than the observed injection-induced response (2.4 ± 1.2 s), likely reflecting the difference in how the mechanical stimulation is applied: proximal (prodding) vs. distal (injection). Direct prodding contact seemed to be required for a robust induction of Ca^2+^ activity, as the ‘unprodded’ muscles were still displaced 23 µm on average but showed little Ca^2+^ activity (**Figure 4D, right**). Prodding elicited an average Ca^2+^ response of 0.6 ΔR/R (**Figure 4D**) while injections elicited an average of 1.17 ΔR/R from non-egg-laying events (**Figure 1D**). This difference in Ca^2+^ activity may result from insufficient activation of both right and left pairs of vulval muscles due the position of the needle along the Z-axis during prodding. Furthermore, vulval muscle prodding did not trigger egg laying, consistent with previous work showing that egg-laying events require coordinated contractions from both anterior and posterior vulval muscles (**Figure 2H**).^28^ Together, these results demonstrate that the vulval muscles are receptive to mechanical stimulation.

**Figure 4:**
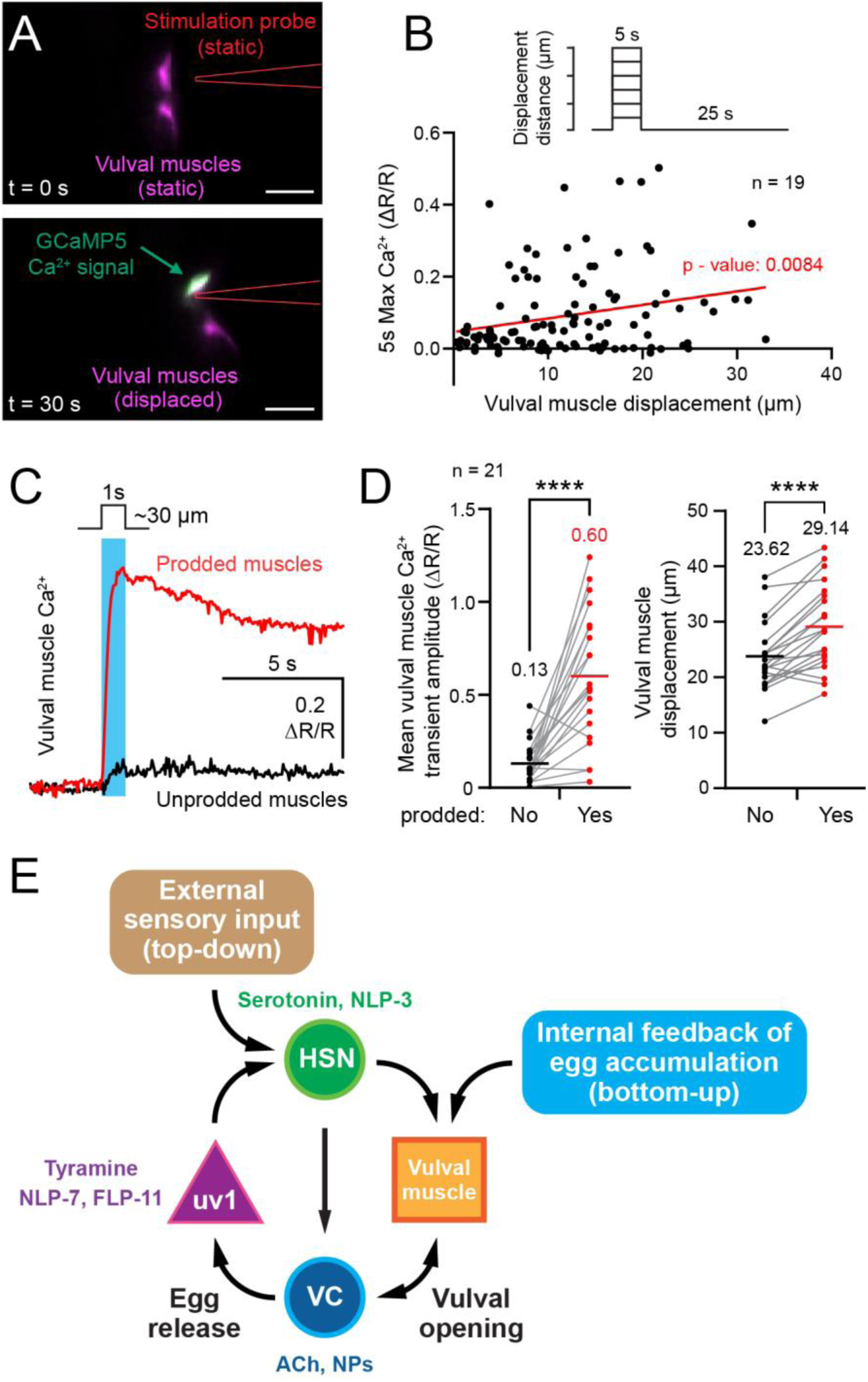
Mechanical stimulation of the vulval muscles elicits a Ca^2+^ response. **(A)** Representative still images of vulval muscle mCherry (magenta) and GCaMP5 (green) fluorescence prior to (top, t = 0 s) or after (bottom, t = 30 s) mechanical stimulation with a stimulation probe (red outline). Scale bar = 25 µm. **(B)** Prodding protocol (top) in which the vulval muscles were prodded for 5 s every 30 s with increasing displacement distances. Scatterplot (bottom) showing the relationship between the determined vulval muscle displacement distance and the max vulval muscle Ca^2+^ response elicited within 5 seconds of the stimulus. Points represent individual prodding responses (n = 114) from 19 animals (p-value = 0.0084, simple linear regression). **(C)** Representative vulval muscle Ca^2+^ traces following a 1 s prodding pulse (vertical cyan line) in unprodded (black, bottom trace) and prodded (red, top trace) vulval muscles. **(D)** Pair-wise comparison (left) of the 5 s mean vulval muscle Ca^2+^ responses in the unprodded (black dots) and their adjacent prodded muscles (red dots). Pair-wise comparison (right) of the calculated vulval muscle displacement for unprodded and prodded muscles. Inset horizontal bars and numbers represent average amplitude and displacement for each group. Asterisks indicate p-value < 0.0001 (paired t-test, n = 21 animals). **(E)** Working model showing how coherent, independent inputs activate the egg-laying cycle. External sensory input into the HSNs (top-down) regulates serotonin and NLP-3 neuropeptide (NP) release. Feedback of egg accumulation by the vulval muscles regulates their activity (bottom-up). Vulval opening mechanically activates acetylcholine (ACh) and NP release from the VC neurons, facilitating muscle contraction, egg release, and activation of the uv1 neuroendocrine cells, restarting the egg-laying cycle.

To test the molecular mechanism mediating the injection-induced vulval muscle response, we investigated the role of PIEZO or TMC mechanosensitive channels. Both channels have been shown to mediate mechanosensitive responses^29–32^ and are expressed in the cells of the egg-laying circuit.^33,34^ However, injection into *pezo-1* (**Figure S4A – C**) or *tmc-1, tmc-2* double mutants (**Figure S4D – F**) triggered robust vulval muscle Ca^2+^ responses with no significant differences in peak amplitude or egg laying when compared to wild-type animals. Overall, our results indicate that while L-type Ca^2+^ channels are required for the injection response (**Figure 1G-I**), Piezo and TMC channels are not. What molecular mechanism mediates the injection mechanosensory response? Assuming the observed kinetics from the GCaMP5 reporter indicate slow (~seconds) response, metabotropic signaling may contribute. Defects in Gα_q_ and Trio RhoGEF signaling block vulval muscle Ca^2+^ activity and egg laying.^35^ Adhesion GPCRs may facilitate mechanosenory responses.^36,37^ In *C. elegans*, latrophilin-1 (LAT-1) mediates mechanosensation in male copulation.^38^ Given this role in a reproductive behavior and LAT-1’s expression in the vulval muscles, HSNs, VCs, and uterine cells,^39^ future work should determine the role of potential adhesion metabotropic mechanosensors in the egg-laying circuit. Alternatively, other cells may mediate a stretch response transmitted to the vulval muscles, leading to depolarization and activation of vulval muscle L-type channels required for Ca^2+^ activity, contractility, and egg laying.^5,35,40^

*C. elegans* body wall muscles (BWMs) have also been shown to respond to mechanical deformations in a manner that does not require *unc-13*, *unc-31*, *pezo-1*, or *tmc-1;tmc-2*.^41^ The BWMs mechanical response is sensitive to amiloride, suggesting DEG/ENaC channels, which are similarly expressed in the vulval muscles^39^ are required for this response.^42,43^ The BWMs do not form direct gap junctions with the egg-laying circuit.^44,45^ They do, however, receive chemical synapses from the VCs, allowing for a possible stretch-sensitive signal from the BWMs to feedback to the VCs and into the vulval muscles, similar to the retrograde signal we described from the vulval muscles to HSNs.^3^ The uterine muscles gap junction onto the vm2 vulval muscles^44^ allowing for possible mechanosensors on the uterine muscles to inform on egg availability. Interestingly, UNC-7 innexins have been shown to function as mechanosensitive hemichannels.^46^ As such, future studies of gap junctions should inform our understanding of how the stretch response is communicated to and between cells of the egg-laying circuit.

We propose a working model for how mechanosensory feedback of stretch modulates the homeostat to drive egg laying in response to injection and in behaving animals (**Figure 4E**). Extensive data has shown that sensory feedback converges onto the HSNs so that eggs are laid in favorable environmental conditions.^47–49^ For example, CO_2_ activates the BAG sensory neurons which release FLP-17 neuropeptides that signal through the EGL-6 receptor on HSN to reduce its activity and inhibit egg laying.^47,49^ Such “top-down” input regulates HSN activity and subsequent release of serotonin and NLP-3 which signal through vulval muscle receptors to drive egg laying.^50^ HSN activity determines ‘when’ and ‘where’ eggs should be laid.^51^ However, “bottom-up” feedback of egg accumulation is still necessary for robust vulval muscle responses to HSN input. Optogenetic stimulation of the HSNs can trigger vulval muscle Ca^2+^ activity in adults,^51^ but it has weaker effects in younger or sterile animals.^3^ Similarly, sterilization of animals with elevated HSN activity significantly reduces vulval muscle Ca^2+^ responses with little effect on HSN activity,^4^ supporting a model in which egg accumulation activates mechanoreceptors to render the vulval muscles sensitive to HSN input.

The ability of the muscles to bypass synaptic input is found in other autonomic reflexes.^52–54^ Decoupling of feedback from these cells to higher control centers does not abolish the reflex but removes the ability of higher control centers to regulate and maintain homeostasis. For example, damage to the mammalian spinal cord above the sacral region leads to loss of faciliatory inputs from the brain, causing an initial suppression of the micturition reflex that is restored over time.^52,53^ A similar pattern occurs upon removal of the HSNs which reduces the frequency of egg-laying active states^5^ and a delay in the onset of egg laying.^3^ The accumulation of a sufficient number of eggs (**Figure 1I**) overcomes deficits in excitatory “top-down” signaling from the HSNs, restoring vulval muscle Ca^2+^ activity^6^ and egg-laying behavior state transitions.^5^ This ‘bottom-up’ processing determines ‘if’ and ‘for how long’ eggs can be laid (**Figure 4E**). These distinct forms and sources of modulatory input help drive optimal patterns of circuit activity that allow animals to adjust their behavior to a changing environment.

## Acknowledgements

We thank Dr. James Baker and members of the Collins Lab for constructive feedback on the manuscript. We thank Drs. Michael Koelle and Ilya Ruvinsky for helpful discussions. We also thank Drs. Xiaofei Bai and Andy Golden for constructing and providing the AG527 strain. We thank Alexander Claman, Addys Bode Hernandez, Robert Fernandez, and Bhavya Ravi for help with strain construction. This work was funded by grants from the National Institutes of Health (NS086932) and the National Science Foundation (IOS-1844657) to KMC. EM was supported by the NSF Graduate Research Fellowship and McKnight Doctoral Fellowship. Some strains were provided by the CGC, which is funded by NIH Office of Research Infrastructure Programs (P40-OD010440).

## Author contributions

Conceptualization: EM and KMC; Methodology: EM; Investigation: EM; Writing: EM and KMC; Funding Acquisition and Administration: EM and KMC.

---

**Figure S1:**
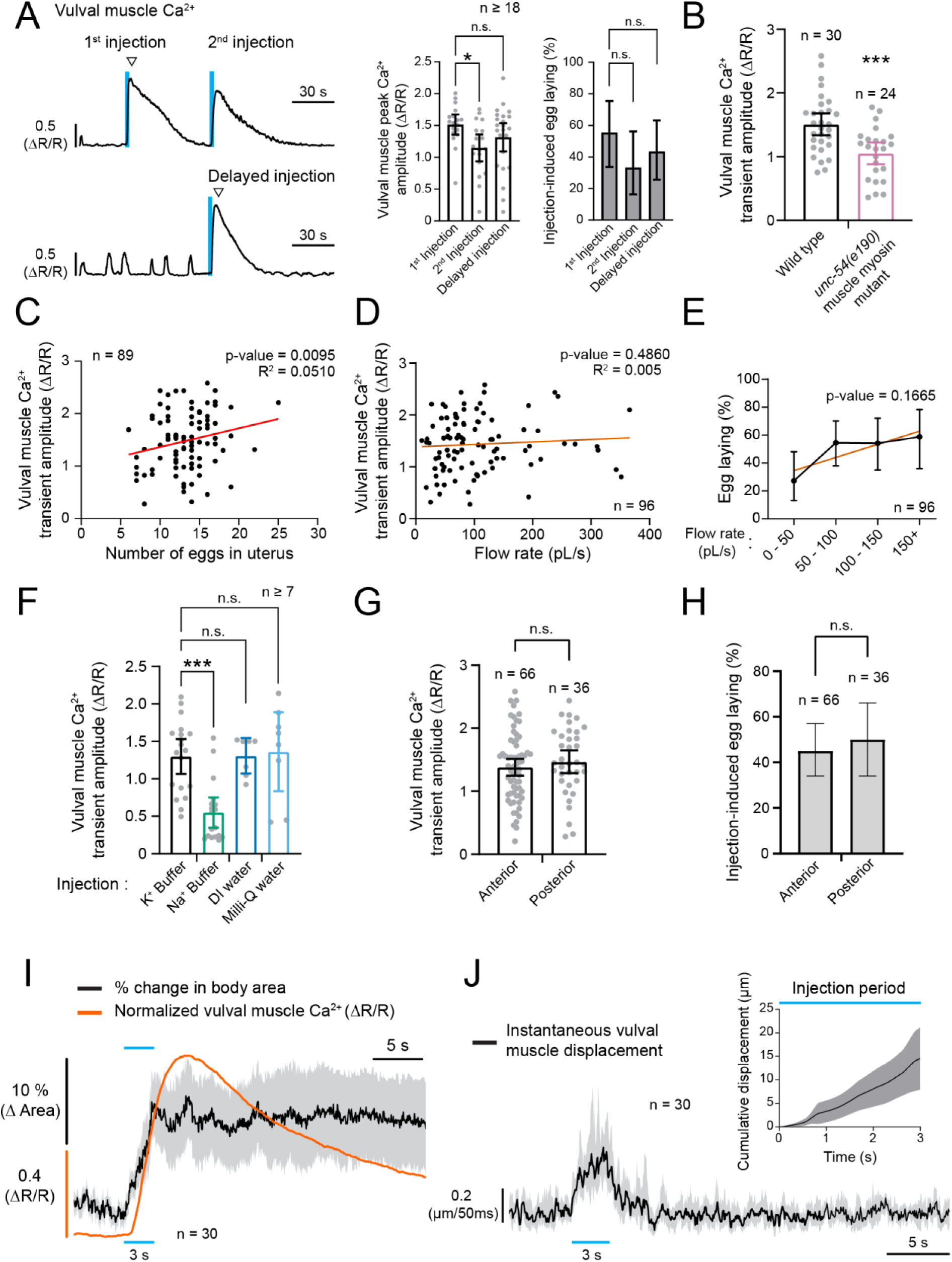
Optimization of the injection protocol. Related to Figure 1. **(A)** Representative Ca^2+^ trace of two, sequential injections into a single animal (top left) compared to a representative Ca^2+^ trace (bottom left) of a single, delayed injection performed at the same time as the second injection on top trace. Vertical cyan lines represent 3 s injection pulses, and trace indicates resulting vulval muscle Ca^2+^ activity. Inverted triangles indicate egg-laying events. Scatterplot with bar graphs (middle) represent average vulval muscle peak Ca^2+^ amplitudes comparing each type of injection (* indicates p-value = 0.0291, ANOVA with Bonferroni’s multiple comparisons test). Solid bar graphs (right) represent percent of injection-induced egg laying in each injection category (n.s. = not significant; Chi-square test). Error bars represent ± 95% confidence interval (C.I.). **(B)** Bar graphs comparing the mean injection-induced vulval muscle Ca^2+^ transient amplitudes following injection into wild-type or *unc-54(e190)* muscle myosin mutant animals (± 95% confidence intervals). Points in scatterplot show responses in individual animals. Asterisks indicate p-value = 0.0004 (unpaired t-test). **(C)** Scatter plot, with a bivariate linear regression comparing the relationship between the number of eggs accumulated inside the uterus at the time of injection and the peak vulval muscle Ca^2+^ response following injection (p-value = 0.0095; standard least squares regression for vulval muscle Ca^2+^ transient amplitude; see ‘statistical analyses’ in STAR methods). **(D)** Scatterplot with linear regression (red line) of relationship between injection flow rate and resulting vulval muscle Ca^2+^ transient amplitude; p-value = 0.4860 (not significant; standard least squares regression for vulval muscle Ca^2+^ transient amplitude; see ‘statistical analyses’ in STAR methods). **(E)** Linear regression of relationship between binned flow rates and percent successful egg laying. p-value = 0.1665 from nominal logistic fit for egg laid (see ‘statistical analyses’ in STAR methods). **(F)** Bar graph with scatterplot showing mean vulval muscle Ca^2+^ transient amplitudes (± 95% confidence intervals) after a 3 s injection with K^+^ buffer (black, n = 19), Na^+^ buffer (green, n = 18), deionized water (blue, n = 7), or Milli-Q water (cyan, n = 8). Asterisks indicate p-value = 0.003; n.s. indicates p-value > 0.05 (Kruskal-Wallis test with Dunn’s correction for multiple comparisons). **(G)** Bar graphs with scatterplot showing average vulval muscle Ca^2+^ transient amplitudes when injections were performed into either the anterior or the posterior gonad. **(H)** Bar graphs with scatterplot showing average percent of animals showing egg release when injections were performed into either the anterior or the posterior gonad. **(I)** Average normalized injection-induced vulval muscle Ca^2+^ activity (orange trace) relative to the average change in whole body area (black trace) following injections into a subsample of 30 wild-type animals. Horizontal cyan bars indicate timing of injection pulse. Grey shaded regions represent ± 95% C.I. **(J)** Average instantaneous vulval muscle displacement in response to injection. Cyan box inset represents the cumulative displacement of the vulval muscles within the 3-second injection period. Shaded regions represent ± 95% C.I.

**Figure S2:**
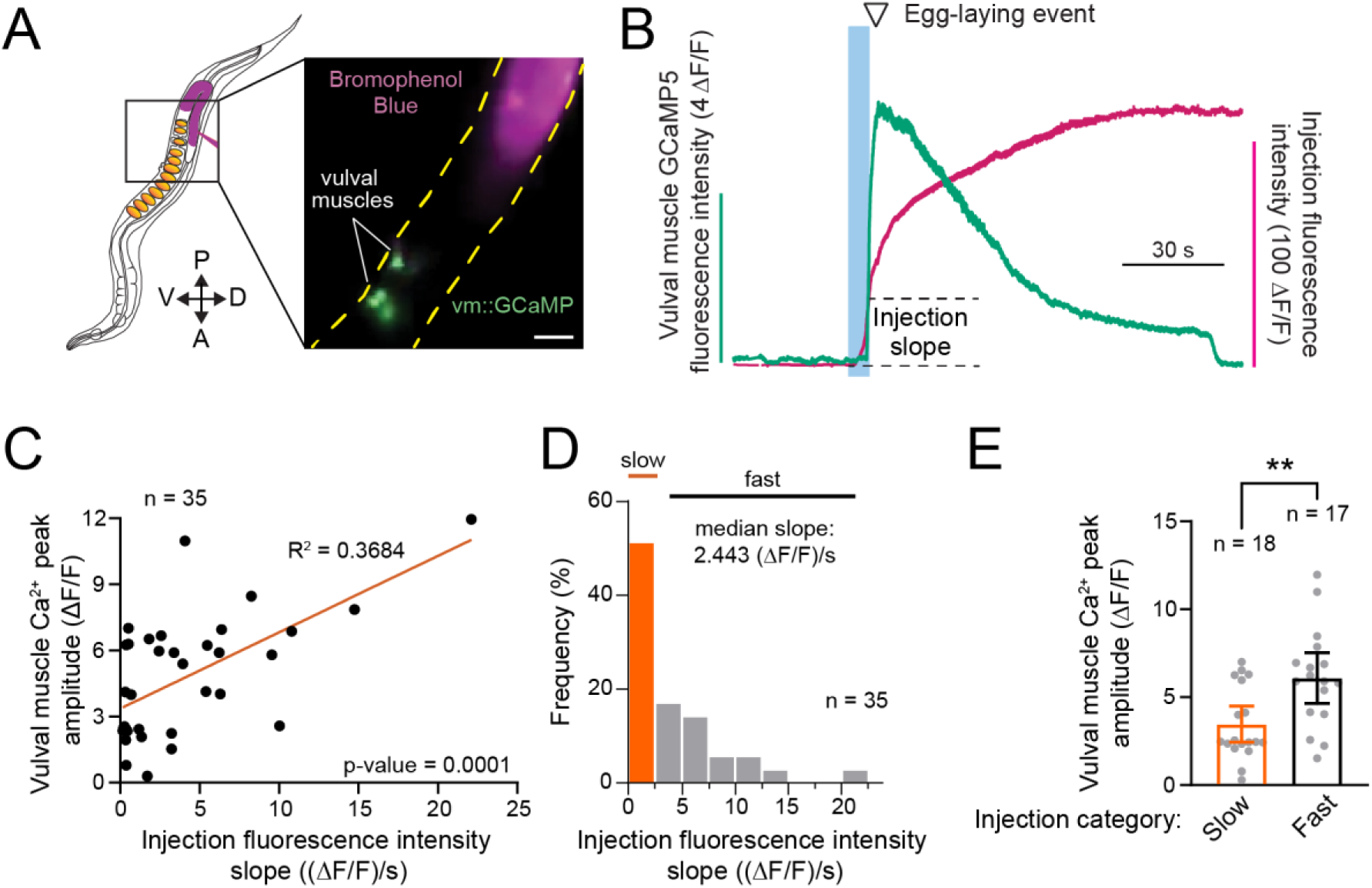
Quantifying correlation of injection flow rate and vulval muscle Ca^2+^ responses using bromophenol blue. Related to Figure 1. **(A)** Schematic of *C. elegans* and micrograph demonstrating injection with bromophenol blue reporter (magenta). Dashed yellow lines represent ventral and dorsal border of animal injected with bromophenol blue (magenta) and the resulted GCaMP5 (green) fluorescence in the vulval muscles (vm). Scale bar = 25 µm. **(B)** Representative traces of vulval muscle GCaMP5 (green; left axis) and bromophenol blue fluorescence (magenta; right axis) after injection used to estimate flow rate and volume delivered. Cyan vertical bar represents time of injection and dotted line represents the change in bromophenol blue fluorescence used to estimate flow rate, e.g. volume injected per 3 s injection time (slope). **(C)** Scatterplot showing significant correlation between injection fluorescence intensity slope (e.g. flow rate) and the resulting vulval muscle Ca^2+^ transient peak amplitude across 35 injected animals (p-value = 0.0001). **(D)** Distribution of all injection fluorescence intensity slopes (flow rates) obtained. Slopes were binned as ‘slow’ (highlighted in orange) or ‘fast’ depending on whether they were below or above the median slope, respectively. **(E)** Bar graphs with scatterplot showing the mean vulval muscle Ca^2+^ transient peak amplitudes seen with ‘slow’ or ‘fast’ injections binned based on slope. Asterisks indicate p-value = 0.0034 (unpaired t-test, n ≥ 17 per group).

**Figure S3:**
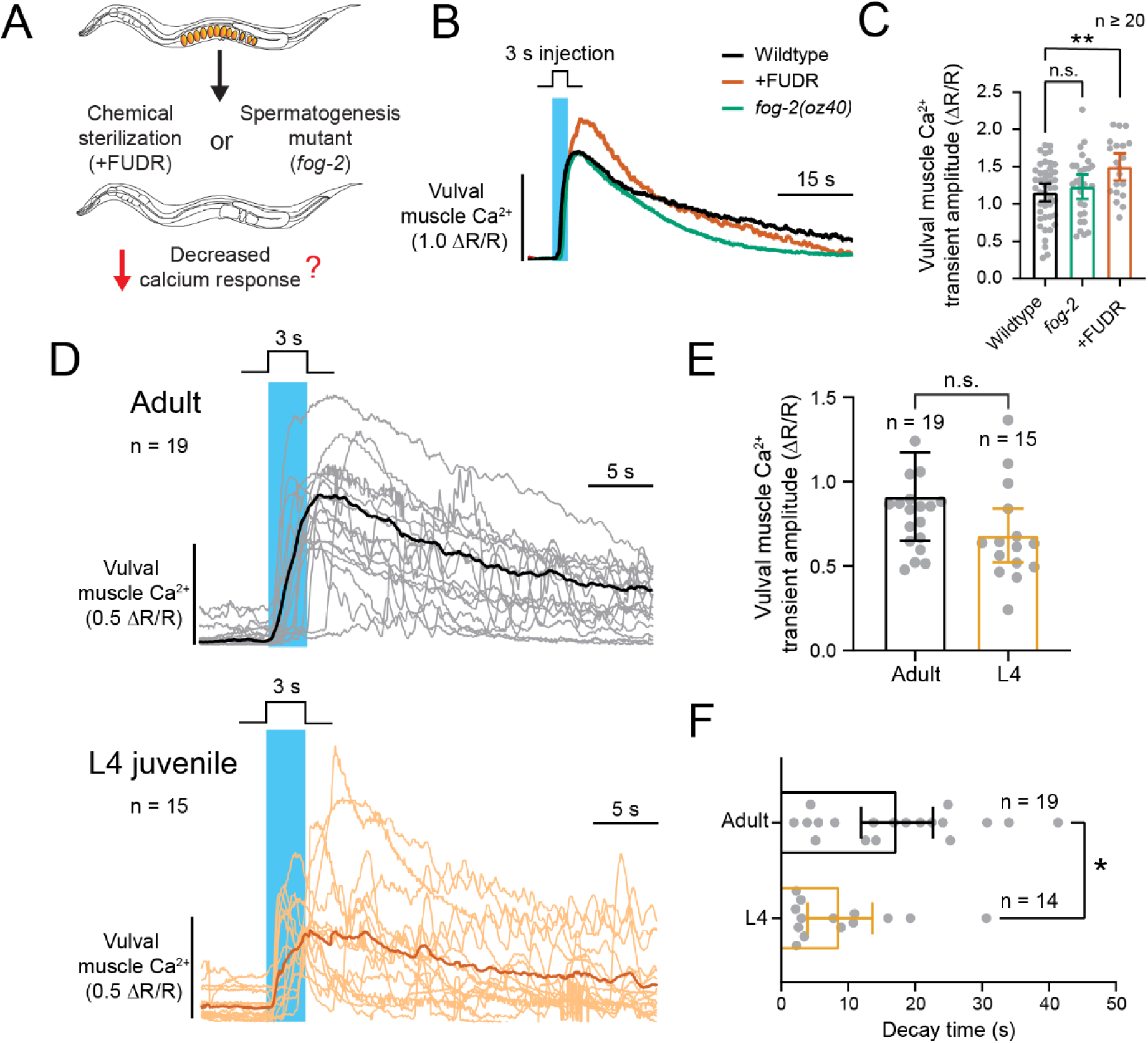
Vulval muscle injection response is independent of egg availability or reproductive maturation. Related to Figure 1. **(A)** Diagram of experimental rationale. Floxuridine (FUDR)-treated wild-type or *fog-2(oz40)* sperm-deficient mutant animals were injected and vulval muscle Ca^2+^ activity was followed using GCaMP5. Orange ovals represent fertilized embryos in the uterus of control animals. **(B)** Representative injection-induced vulval muscle Ca^2+^ responses of control wild-type (black), wild-type sterilized with FUDR (red), and *fog-2(oz40)* mutant animals (green). **(C)** Bar graphs with scatterplot showing mean vulval muscle Ca^2+^ transient amplitudes following microinjection. Asterisk indicates p-value = 0.0044, and n.s. indicates p-value > 0.05 (one-way ANOVA with Bonferroni’s multiple comparisons test; n ≥ 20 animals per condition). **(D)** Injection-induced Ca^2+^ responses for 1-day old adult (top; black trace) and late-L4 (bottom; orange trace) animals. Solid lines show the average Ca^2+^ responses, and lighter lines show responses from individual animals (adults, n = 19; L4 juveniles, n = 15). **(E)** Bar graphs with scatterplots showing mean vulval muscle Ca^2+^ amplitudes. n.s. indicates p-value > 0.05 (Mann-Whitney test). **(F)** Bar graphs with scatterplots showing mean decay time. Asterisk indicates p-value = 0.0139. Vertical cyan bars show timing of 3 s microinjection. To facilitate visual comparison, one data point with vulval muscle Ca^2+^ transient amplitude value of 2.99 (ΔR/R) from an adult animal is not shown in D and E.

**Figure S4:**
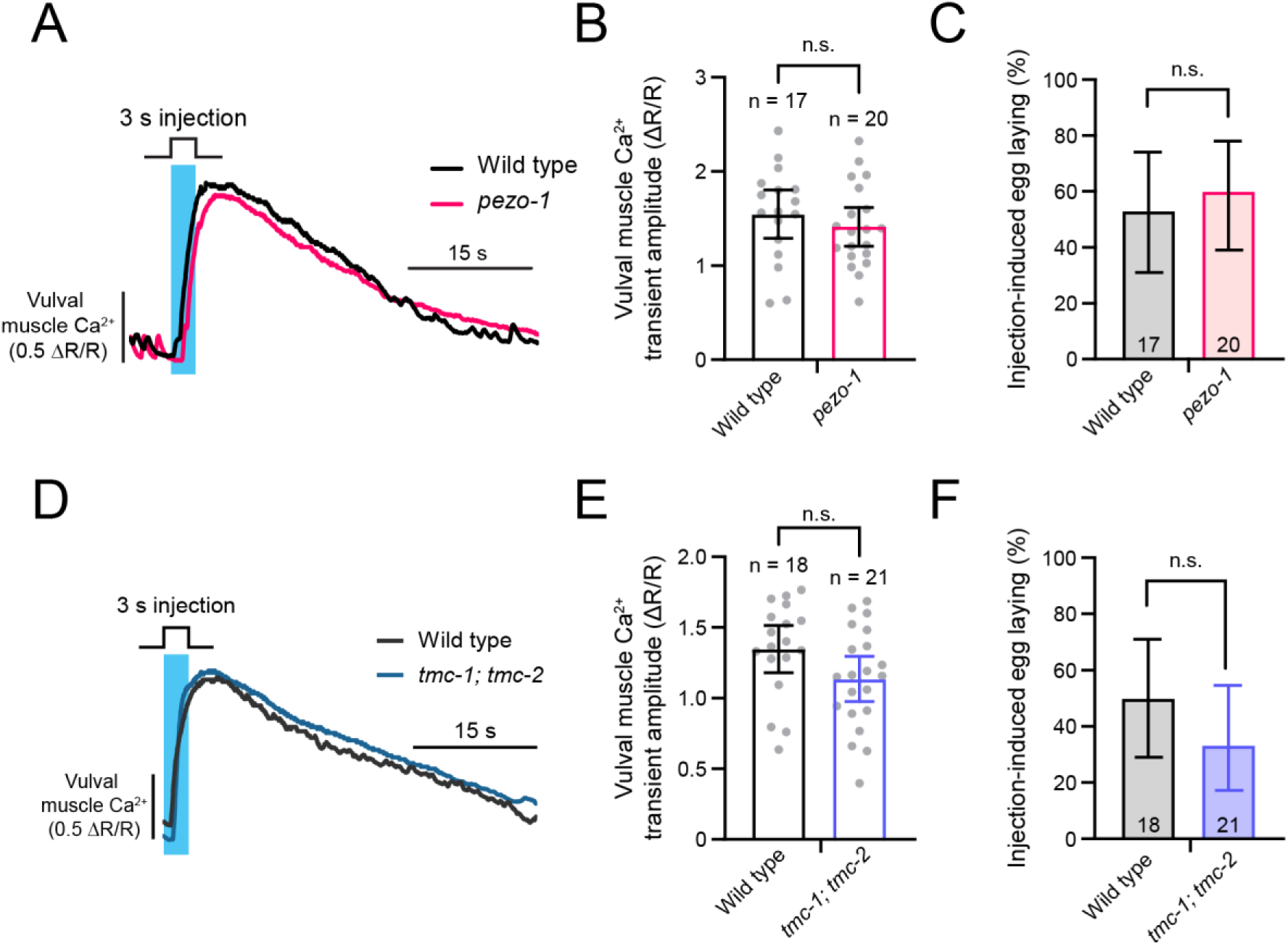
PIEZO and TMC channels are not required for the injection-induced vulval muscle Ca^2+^ and egg-laying response. Related to Figure 4. **(A)** Representative injection-induced vulval muscle Ca^2+^ traces after injection into wild-type (black) and *pezo-1(av149)* mutant animals (magenta). (B) Bar graphs with scatterplot showing mean vulval muscle Ca^2+^ responses of wild-type (black, n = 17) and *pezo-1* mutants (magenta, n = 20). Error bars indicate 95% confidence interval; n.s. indicates p-value > 0.05 (unpaired t-test). (C) Bar graphs showing percent of injections that resulted in an egg-laying event (mean ± 95% confidence intervals for the proportion); n.s. indicates p-value > 0.05 (Fisher’s exact test). (D) Representative injection-induced vulval muscle Ca^2+^ traces after injection into wild-type (black) and *tmc-1(ok1859); tmc-2(ok1302)* double mutant animals (blue). (E) Bar graphs with scatterplot showing mean vulval muscle Ca^2+^ responses of wild-type (black, n = 18) and *tmc-1; tmc-2* double mutants (blue, n = 21). Error bars indicate 95% confidence interval for the mean; n.s. indicates p-value > 0.05 (unpaired t-test). (F) Bar graphs showing percent of injections that resulted in an egg-laying event (mean ± 95% confidence intervals for the proportion); n.s. indicates p-value > 0.05 (Fisher’s exact test). Vertical cyan bars indicate 3 s injection. Numbers inside bar graphs indicate sample sizes.

## STAR Methods

### RESOURCE AVAILABILITY

#### Lead contact

Further information and requests for resources and reagents should be directed to and will be fulfilled by the lead contact, Kevin M. Collins.

#### Materials availability

Plasmids and strains generated in this study are available from the lead contact by request.

#### Data and code availability

- Microscopy, ratiometric traces, and summary data analyzed for statistical significance reported in this paper will be shared by the lead contact upon request.
- This paper does not report original code.
- Any additional information required to reanalyze the data reported in this paper is available from the lead contact upon request.

### EXPERIMENTAL MODEL AND SUBJECT DETAILS

Experiments were performed using adult *Caenorhabditis elegans* N2 hermaphrodites staged 24 h past the late L4 larval stage. For assays involving juvenile animals, fourth-larval stage animals were picked 30 minutes prior to the assay. *Caenorhabditis elegans* strains were grown at 20°C on Nematode Growth Medium (NGM) agar plates with *E. coli* OP50 grown in B broth as a food source.^57^ A list of strains, mutants, and transgenes used in this study can be found in the **Key Resources Table.**

### METHOD DETAILS

#### Calcium reporter transgenes and strain construction

##### Vulval muscle Ca^2+^

Vulval muscle Ca^2+^ activity was recorded in adult animals using LX1918 *vsIs164 [unc-103e::GCaMP5::unc-54 3’UTR + unc-103e::mCherry::unc-54 3’UTR + lin-15(+)]*, *lite-1(ce314), lin-15(n765ts) X*, as described.^6^ Because the *unc-103e* promoter does not express strongly in the vulval muscles of younger L4 animals,^3^ MIA51 *keyIs12 [ceh-24::GCaMP5::unc-54 3’UTR ceh-24::mCherry::unc-54 3’UTR lin-15(+)] IV, lite-1(ce314), lin-15(n765ts) X*, was used in experiments comparing injection responses in late L4 animals as described.^3^

To image vulval muscle Ca^2+^ activity in animals defective for muscular contractions, MIA206 [*unc-54(e190) I; vsIs164 X lite-1(ce314) lin-15(n765ts) X*] strain was generated. To generate this strain, LX1918 males were crossed with CB190 [*unc-54(e190) I*] myosin heavy chain null mutant hermaphrodites.^57,58^ Offspring were allowed to self and uncoordinated, mCherry(+) F2 progeny were then selected and allowed to self for multiple generations until both phenotypes were observed to be homozygous.

To record Ca^2+^ activity from the vulval muscles of animals lacking fertilized embryos, BS553 [*fog-2(oz40) V)*] spermatogenesis mutant^17^ males were crossed with LX1918 hermaphrodites. Cross-progeny hermaphrodites were picked and were crossed back to BS553 *fog-2* males to generate mCherry(+), *fog-2/+* males and mCherry(+), *fog-2* hermaphrodites which were identified from single mCherry(+) L4 hermaphrodites that then developed into sterile, *fog-2* adults. Male and hermaphrodite progeny continued to be crossed until the *fog-2*, mCherry(+) phenotypes were homozygous. The resulting strain, MIA34 [*fog-2(oz40) V; lite-1(ce314) lin-15(n765) vsIs164 X*], was then maintained by male and female self-progeny.

To determine if injection-induced vulval muscle Ca^2+^ responses were dependent on input from the HSNs or VC neurons, a strain lacking HSNs while simultaneously expressing Tetanus Toxin in the VC neurons was generated. To do this, N2 males were crossed with the HSN-deficient LX1938 *egl-1(n986dm) V; lite-1(ce314) vsIs164 lin-15(n765ts) X* hermaphrodites.^6^ Resulting male cross-progeny were then mated with MIA144 *keyIs33; lite-1(ce314) lin15(n765ts) X* hermaphrodites^19^ expressing Tetanus Toxin in the VC neurons from a minimal *lin-11* enhancer^59^ which imparts a mild uncoordinated reversal phenotype. Resulting cross-progeny L4s were then picked, allowed to self, and were then selected for mCherry(+), Egl, and Unc reversal phenotypes. Animals were then positively genotyped for *egl-1(n986dm)* using the ‘egl-1_fwd’ and ‘egl-1_rev’ oligos (**Key Resources Table**) followed by NciI enzyme digestion. Confirmed homozygous animals were kept as strain MIA289 *keyIs33; egl-1(n986dm) V; vsIs164 lite-1(ce314) lin-15(n765ts) X*.

To confirm the lack of development of the vulval muscles in *egl-17* sex myoblast migration mutants, MT3177 *egl-17(n1377),*^18^ hermaphrodites were crossed with vsIs164 mCherry(+) males. Progeny hermaphrodites were allowed to self and then selected for mCherry(+) and Egl phenotypes until homozygous. The resulting MIA555 [*egl-17(n1377) vsIs164 lite-1(ce314) lin-15(n765ts) X*] strain was then subjected to fluorescence imaging to confirm lack of proper vulval muscle development (**Figure 2G**).

To eliminate neuropeptide signaling while monitoring injection-induced Ca^2+^ activity in the vulval muscles, the dense core vesicle release mutant, CB928 *unc-31(e928) IV*,^57^ was crossed with LX1918 males. *vsIs164* mCherry(+) hermaphrodite cross-progeny were allowed to self and were then selected for mCherry(+), Unc, and Egl phenotypes, resulting in strain MIA312 [*unc-31(e928) IV; vsIs164 X; lite-1(ce314) lin-15(n765ts) X*]. To further investigate the role of neuropeptide signaling, two strains defective in neuropeptide synthesis, VC671 *egl-3(ok979) IV* and MT1071 *egl-21(n476) IV*,^23,24^ were crossed with LX1918 males. Resulting cross progeny were selected for mCherry(+) and Egl phenotypes and allowed to self until homozygous. This generated MIA543 [*egl-3(ok979) V; vsIs164; lite-1(ce314) lin-15(n765ts) X*] and MIA544 [*egl-21(n476) IV; vsIs164 lite-1(ce314) lin-15(n765ts) X*] strains.

Similarly, to eliminate neurotransmitter release while monitoring injection-induced Ca^2+^ activity from the vulval muscles, the neurotransmitter release mutant, EG9631 *unc-13(s69) I*,^60^ was crossed with *vsIs164* mCherry(+) males. mCherry(+) hermaphrodite cross-progeny were picked as L4s, allowed to self, and mCherry(+), Unc F2 progeny were kept, resulting in strain MIA316 *unc-13(s69) I; vsIs164 lite-1(ce314) lin-15(n765ts) X*. To eliminate potential tyramine signaling from the uv1s or other cells, LX1996 [*tdc-1(n3419) II; vsIs164 lite-1(ce314) lin-15(n765ts) X*] animals lacking tyramine biosynthesis and expressing GCaMP5 and mCherry in the vulval muscles, were used. To generate this strain, LX1836 males,^6^ expressing Channelrhodopsin-2 in the HSNs, were mated with LX1932 hermaphrodites, expressing Channelrhodopsin-2 in the HSNs and GCaMP5 and mCherry in the vulval muscles.^6^ The resulting male progeny were then mated with MT13113 *tdc-1(n3419) II* tyramine synthesis mutant^25^ hermaphrodites. GFP(+) hermaphrodite cross-progeny were then allowed to self and were genotyped for the *tdc-1* mutation using the ‘tdc-1-MF’, ‘tdc-1-WTF’, and ‘tdc-1-WTR’ oligos (**Key Resources Table**). Absence of the Channelrhodopsin-2 transgene was confirmed via absence of the fluorescent reporter.

To image Ca^2+^ activity in the vulval muscles of animals lacking the EGL-19, L-type voltage-gated Ca^2+^ channel,^27^ LX1918 males were crossed with MT1212 *egl-19(n582) IV* hermaphrodites. Cross-progeny were isolated and allowed to self and resulting mCherry(+) non-Egl progeny were picked and allowed to self until homozygous for mCherry(+) and Egl phenotypes. The resulting strain MIA185 [*egl-19(n582) IV; lite-1(ce314) lin-15(n765ts) vsIs164 X*] was used in this study.

To monitor Ca^2+^ activity in the vulval muscles of mechanosensory mutants, *vsIs164* males were crossed with *pezo-1* mutants provided by Drs. Xiaofei Bai and Andy Golden.^33^ This resulted in the generation of the AG527 strain [*pezo-1 (av149) IV; vsIs164 lite-1(ce314) lin-15(n765ts) X*].

To visualize vulval muscle Ca^2+^ activity in tmc-1, tmc-2 mechanosensory mutants, N2 males were crossed with LX1919 vsIs165 [unc-103e::GCaMP5::unc-54 3’UTR + unc-103e::mCherry::unc-54 3’UTR + lin-15(+); lite-1(ce314) lin-15(n765ts) X] hermaphrodites.^4,6^ mCherry(+) vsIs165/+ male cross-progeny were then crossed with RB1546 tmc-1(ok1859) X hermaphrodites. mCherry(+) cross-progeny hermaphrodites were then picked and allowed to self. mCherry(+) cross-progeny were then genotyped for tmc-1(ok1859) using a duplex PCR strategy with the following primers:^61^ ‘tmc-1(ok1859) outside_Fwd’, ‘tmc-1(ok1859) inside_Rev’, and ‘tmc-1(ok1859) outside_Rev’ (**Key Resources Table**), generating MIA486 vsIs165; tmc-1(ok1859) X strain used in this study. To generate MIA487 vsIs165 tmc-2(ok1302) X, the same approach was used but the RB1237 tmc-2(ok1302) X hermaphrodites was used instead when crossed with vsIs165/+ males. mCherry(+) progeny were confirmed to be homozygous for tmc-2(ok1302) using the following primers: ‘tmc-2(ok1302) outside_Fwd’, ‘tmc-2(ok1302) inside_Fwd’, and ‘tmc-2(ok1302) outside_Rev’ (**Key Resources Table)**. During construction of MIA486 and MIA487, we noticed the loss of the lite-1(ce314) X mutation present in the original LX1919 starting strain as tmc-1 and tmc-2 genes, both of which are near lite-1, were being homozygosed. To account for any changes in vulval muscle Ca^2+^ activity that arise from blue light illumination rather than acute injection, a control strain MIA485 vsIs165 was generated by crossing N2 males with LX1919 hermaphrodites. Heterozygous cross-progeny hermaphrodites with mCherry expression were selected, selfed, and loss of lite-1(ce314) in the progeny was confirmed through genotyping.

MIA494 *vsIs165*; *tmc-1(ok1859) tmc-2(ok1302) X* was generated to image vulval muscle Ca^2+^ activity in animals lacking both TMC-1 and TMC-2. Briefly, N2 males were mated to RB1546 *tmc-1(ok1959)* hermaphrodites, and the resulting hemizygous (*tmc-1/Ø*) males were then crossed with MIA487 hermaphrodites. The resulting cross-progeny were then allowed to self until *tmc-1* and *tmc-2* mutations were confirmed to be homozygous through genotyping.

##### uv1 neuroendocrine cell Ca^2+^

Ca^2+^ activity of uv1 neuroendocrine cells was monitored in adult animals using MIA136 *keyIs34 lite-1(ce314) lin-15(n765ts) X* which expresses GCaMP5 and mCherry from the *tdc-1* promoter^25^ which provides stronger uv1 expression compared to the *ocr-2* promoter previously used.^6^ To generate this strain, the *ocr-2* promoter was replaced with a 1.5 Kb region of the *tdc-1* promoter amplified from *Ptdc-1::ChR2* generously provided by Mark Alkema using oligonucleotides ‘Ptdc-1new-fwd’ and ‘Ptdc-1-new-rev’ (**Key Resources Table**)digested with SphI/XmaI, and ligated into similarly digested pKMC281 [*Pocr-2::mCherry::ocr-2(3’UTR)*] or pKMC284 [*Pocr-2::GCaMP5::ocr-2(3’UTR)*], generating pAB5 [*Ptdc-1::mCherry::ocr-2(3’UTR)*] and pAB6 [*Ptdc-1::GCaMP5::ocr-2(3’UTR)*]. pAB5 [*Ptdc-1::mCherry* (5 ng/µL)], pAB6 [*Ptdc-1::GCaMP5* (20 ng/µL)], and pL15EK (50 ng/µL)^62^ were injected into LX1832 *lite-1(ce314), lin-15(n765ts)* animals, generating the extrachromosomal transgene *keyEx28* which demonstrated bright and specific expression in the uv1 cells along with dim expression in the RIM neurons. This transgene was then integrated to chromosomes using UV/TMP,^63^ generating four independent integrants *keyIs34–37*, of which *keyIs34* was used for this study.

To determine the injection-induced response of the uv1s in animals lacking proper vulval muscle development, the MIA546 [*egl-17(n1377) keyIs34 lite-1(ce314) lin-15(n765ts) X*] strain was generated. MT3188 *egl-17(n1377)* males were crossed with MIA136 hermaphrodites. Cross-progeny were then pooled into multiple plates, allowed to self, and mCherry(+), Egl progeny were then allowed to self until both were confirmed visually to be homozygous.

The MIA332 [*unc-54(e190) I; keyIs48; keyIs34 lite-1(ce314) lin-15(n765ts) X*] strain lacking proper vulval muscle contraction was generated to test if contractility is required for the injection-induced uv1 Ca^2+^ response. To generate this strain, MIA239 males (expressing Channelrhodopsin-2 in the vulval muscles along with mCherry and GCaMP5 in the uv1s) were crossed with MIA298 *unc-54(e190)* hermaphrodites expressing Channelrhodopsin-2 in the vulval muscles along with GCaMP5 and mCherry in the VC neurons.^19^ Cross progeny were selected for mCherry(+) in the uv1 cells but not in the VC neurons. Thereafter, Unc and mCherry(+) progeny were selected until homozygous.

##### VC Ca^2+^

Ca^2+^activity in the VC neurons was monitored in adult animals using strain LX1960 *vsIs172; lite-1(ce314) lin-15(n765ts) X,* as described.^6^ Ca^2+^ activity levels were averaged from all VC cell bodies and process (VC1-VC6). However, due to expression patterns, differences in focus, and the position of worms during recording, not all VCs were captured in all experiments, especially VC1 and VC6 which are more distant from the vulval-proximal VC4 and VC5.

To determine the injection-induced response of VCs in animals lacking proper vulval muscle development, MT3188 *egl-17* males were crossed with LX1960 hermaphrodites. Cross-progeny were then pooled into multiple plates, allowed to self, and cross progeny were identified via mCherry(+) and Egl phenotypes. Animals were then singly picked and allowed to self until homozygous, resulting in the MIA547 strain [*vsIs172; egl-17(n1377) lite-1(ce314) lin-15(n765ts) X*].

To test the role of muscle contractility on the injection-induced VC response, N2 males were crossed with LX1960 hermaphrodites. Cross-progeny males were then crossed with CB190 hermaphrodites. F1 mCherry(+) progeny were selected and subsequent generations were allowed to self until homozygous for uncoordinated phenotype and mCherry(+), resulting in the MIA534 strain [*unc-54(e190) I*; *vsIs172; lite-1(ce314) lin-15(n765ts) X*].

##### HSN Ca^2+^

Ca^2+^ activity in the HSNs was monitored in adult animals using LX2004 *vsIs183 lite-1(ce314) lin-15(n765ts) X,* as described.^6,64^ To determine the role of the vulval muscles in the injection-induced HSN Ca^2+^ response, we generated the MIA545 strain [*egl-17(n1377) vsIs183 lite-1(ce314) lin-15(n765ts) X*]. To do this, MT3188 *egl-17* males were crossed with LX2004 hermaphrodites. Progeny were pooled, allowed to self, and cross-progeny were identified via mCherry(+) and Egl phenotypes. These animals were cloned to plates singly and allowed to self over multiple generations until homozygous.

To image Ca^2+^ dynamics in the HSNs of vulval muscle contractility mutants, we used the MIA205 strain [*unc-54(e190) I; vsIs183 lite-1(ce314) lin-15(n765ts) X*]. Briefly, LX2004 males were crossed with CB190 *unc-54* mutant hermaphrodites. Non-unc, mCherry(+) progeny, were picked, allowed to self, and homozygous mCherry(+) Unc progeny were kept.

#### Egg accumulation at onset of egg laying

To determine the extent of egg accumulation at onset of egg laying, both wild-type and MIA26 *egl-1(n986dm)* mutant animals were staged following the L4 molt as previously described.^3^ Animals were observed for the first instance of egg laying every 30 minutes (after a 4 h period for wild type and after a 16 h period for *egl-1(dm)* mutants). Once an animal had laid its first egg, it was dissolved in a 20% bleach solution, and the eggs in the uterus were counted as described.^65^ The number of eggs in the uterus plus the number of eggs laid on the plate were considered the total number of eggs accumulated at the onset of egg laying.

#### Microinjection protocol and flow rate estimate

To induce acute increases in pressure within *C. elegans* worms, we performed microinjections essentially as described.^7^ To create microinjection needles, filamented glass capillaries (4”, 1.0 mm outer diameter; World Precision Instruments, 1B100F-4) were pulled using a Sutter P-97 Flaming/Brown type micropipette puller using a custom program with the following parameters: Pressure = 500, Heat = 632 (ramp test value), Pull = 45, Velocity = 75, Delay = 90. Needles were back-filled with injection buffer (2% polyethylene glycol, 20 mM potassium phosphate, 3 mM potassium citrate, pH 7.5) and then placed into a Narishige HI-7 injection holder. Worms were injected using a Narishige IM-300 pneumatic microinjector with injection pressure between 30 – 40 psi, unless otherwise noted. Injections were programmed for a 3 s injection pulse and initiated manually using a foot pedal. To synchronize injection onset with captured brightfield and fluorescence micrographs, the ‘sync’ voltage signals of the microinjector were controlled and/or monitored using an Arduino Uno microcontroller running the Firmata library in Bonsai.^66^ Injections (3 s) were triggered either using a foot-pedal or a TTL digital pin, and the timing of injection onset was recorded using an analog pin. To assess consistency of injection needles, the average flow rate of needles was calculated from brightfield recordings by measuring the volume of a spherical drop of injection buffer injected into halocarbon oil. The volume was calculated by measuring the expanding diameter of the drop in the first 20 frames of a three second injection (10 msec exposures recorded at 20 Hz). The average flow rate of multiple needles pulled using this protocol was calculated to be 47 pL/s with a standard deviation of 15 pL/s (n = 10).

To perform microinjections, worms were placed into a small drop of halocarbon oil on a 2% agarose pad to immobilize them.^7^ Animals were then placed onto an inverted Zeiss Axio Observer.Z1 fluorescence microscope for imaging. A Narishige coarse manipulator (model MMN-1) and a Narishige three-axis oil hydraulic micromanipulator (model MMO-203) were used to position the tip of the needle next to the gonad of the worm. Needles were then introduced along the dorsal surface into either anterior or posterior gonad of the worm before initiating the recording (see fluorescence imaging). The first 3 s injection was triggered 30 seconds after the onset of synchronous brightfield and fluorescence recording. Subsequent injections were performed in one-minute intervals. Before the end of the recording, the injection needle was withdrawn from each animal, and the flow rate of the needle was once again recorded to determine if the flow rate had changed materially during the injection. The median flow rate from a subsample of 104 animal injections was 82 pL/s and ranged between 10 and 365 pL/s. Such variation was seen to result from either needle tips clogging or being open further as a consequence of injection, but this variability did not seem to alter injection-induced responses (see optimization of microinjection protocol).

#### Brightfield and fluorescence recordings

To record egg-laying behavior during injections, a Grasshopper 3 camera (FLIR) was used to capture 2 x 2 binned, 1,024 x 1,024 8-bit JPEG image sequences. Vulval muscle, HSN, VC, or uv1 Ca^2+^ responses were recorded as described.^6,64^ GCaMP5 and mCherry fluorescence was excited at 470 nm and 590 nm, respectively, for 10 msec using a Colibri.2 LED illumination system. Recordings were collected at 20 fps and a 256×256 pixel resolution (4×4 binning) using a Hamamatsu Flash 4.0 V2 sCMOS camera recording at 16-bit depth through a 20x Apochromatic objective (0.8NA). GCaMP5 and mCherry channels were separated via a Gemini beam-splitter. Worms were recorded for 30 seconds prior to injection to establish baseline levels of cell activity and were completed within five minutes to reduce effects from worm desiccation after prolonged exposure to halocarbon oil. Image sequences were imported into Volocity (Version 6.5.1, Quorum Technologies Inc.) for GCaMP5/mCherry ratiometric analysis, image segmentation, Ca^2+^ quantitation of GCaMP5/mCherry ratio changes (ΔR/R), and Ca^2+^ transient peak finding, as described previously.^6,64^

#### Optimization of microinjection protocol

To ensure injections into control animals and the different mutants were comparable, the correlation between flow rate and recorded calcium transient amplitudes was analyzed. From this comparison, we found that flow rate had no significant effect on the amplitude of the resulting vulval muscle Ca^2+^ transients (p-value = 0.4860; **Figure S1C**) and, as a result, the injection flow rate was allowed to vary. Additionally, the correlation between flow rate and an egg-laying response was also analyzed. This relationship also showed no significant correlation between flow rate and percent of injection-induced egg laying (p-value = 0.1665; **Figure S1E**. Individual animals were injected twice to determine if subsequent responses were comparable. We found that subsequent injections resulted in lower Ca^2+^ transient amplitudes and were but equally likely to result in an egg-laying event (**Figure S1A**) However, given the wide range of flow rates obtained among multiple injections, we limited our analyses of injection-induced Ca^2+^ responses to each animal’s first injection.

To determine if the composition of the injection buffer influenced the Ca^2+^ response, worms were injected with deionized water or Milli-Q water. Injections with both types of water did not differ in their ability to induce comparable levels of Ca^2+^ transient activity from the vulval muscles (p-value >0.05; **Figure S1F**). However, we observed that when the injection buffer was made with Na^+^ salts instead of K^+^ salts, a decrease in the average vulval muscle Ca^2+^ response was observed (p-value = 0.0003; **Figure S1F**) when compared to the standard K^+^ salt containing buffer. To ensure that the injection-induced response is not a consequence of a disruption of the Na^+^/K^+^ gradient, low flow rate injections were performed by decreasing injection pressure to 20 psi. In these low-flow experiments, we observed that injections were more likely to fail to elicit a Ca^2+^ response even when the ‘optimal’ K^+^ buffer was used. Injections were performed into either the anterior or posterior gonad, with no significant difference in vulval muscle Ca^2+^ transient amplitude or proportion of injection-induced egg laying observed (**Figure S1G-H**). Attempts to determine if injections performed in other compartments (e.g. the uterus) also drove vulval muscle Ca^2+^ activity were unsuccessful due to injection needles frequently getting clogged (data not shown).

##### Injections with bromophenol blue

To visualize injections, and to determine if Ca^2+^ dynamics correlated with the rate of liquid spread at lower flow rates (injection pressure 25 psi), a solution of 450 µM bromophenol blue made in injection buffer was used as a fluorescent tracker (**Figure S2A**). Bromophenol blue was excited at 590 nm alongside co-expressed mCherry and collected through the same mCherry filter set. To determine if the flow rate of injections influenced vulval muscle Ca^2+^ responses, worms were manually injected (25 psi) by depression of the foot pedal until the bromophenol fluorescence was seen by eye to reach the vulval muscles (**Figure S2A**). Bromophenol fluorescence was then quantified in Volocity. The Sum of individual pixel intensities was used as a proxy of total volume delivered during the course of each injection [(pixel area) x (16-bit fluorescence intensity for each pixel above baseline) and with all such pixels summed]. Changes in bromophenol fluorescence intensity and area over time, after background subtraction, were used to calculate the ΔF/F.

To determine total volume injected, the maximum fluorescence intensity of all injections within a 2-minute period after injection was determined and used as a proxy for volume injected. The flow rate of each injection was calculated by measuring the slope of the injection fluorescence intensity during the injection period (**Figure S2B**). At the same time, GCaMP5 fluorescence intensity (ΔF/F) from the vulval muscle response was quantified. From this, a significant correlation was observed between the slope of the bromophenol blue fluorescence intensity during injection and the vulval muscle GCaMP5 fluorescence peak amplitude (**Figure S2C**). Injections were then binned into two groups depending on the median value of all slopes obtained. All calculated slopes lower than the median were categorized as a ‘slow’ injection while injections above the median were categorized as ‘fast’ injections (**Figure S2D**). From this, ‘slow’ injections were seen to elicit lower peak Ca^2+^ amplitudes when compared to ‘fast’ injections (**Figure S2E**). This indicated that the rate at which an injection was performed significantly determined the strength of the resulting vulval muscle Ca^2+^ response but only for injections performed at these lower-than-typical injection pressures (25 psi). Since flow rate did not significantly correlate with Ca^2+^ dynamics at injection pressures greater than 30 psi, to optimize Ca^2+^ responses, and to reduce instances of needles clogging, injection pressures used when testing animal responses were maintained between 30 – 40 psi.

#### Injection-induced increase in area measurements

To characterize the physical increase in body size observed following injections, a subsample of 30 animals was chosen at random and area measurements were performed. To do this, brightfield recordings of injections were thresholded using ImageJ and imported into Volocity for automated object finding and area measurements. The relative change in area was calculated per individual and pooled to generate the average change in area following injections (**Figure S1I**).

#### Vulval muscle prodding protocol

To determine if the vulval muscles were receptive to mechanical stimulus, worms were immobilized as per the microinjection protocol. A prodding needle with a 2.7 µm wide tip was positioned next to either the anterior or posterior part of the vulva. To drive a mechanical displacement onto the prodded vulva, a motorized stage was moved at a range of distances (10 – 60 µm) in the direction of the needle to determine the optimum displacement distance. Subsequent experiments used a blunt prodding probe (4.6 µm wide tip), displaced 50 µm for one second. Image sequences were then segmented into paired prodded and unprodded vulval muscle recordings as per the fluorescence recording protocol. Paired Ca^2+^ responses in the 5 s period following the prodding stimulus were averaged and analyzed.

#### Vulval muscle displacement measurements

##### Vulval muscle displacement via injection

To determine if microinjections led to physical movement of the vulval muscles, the instantaneous displacement was calculated for the random subsample of animals used to calculated average change in body area indicated above. To determine displacement, the vulval muscles centroid position was determined in Volocity. Displacement was calculated by measuring instantaneous changes in centroid X and Y position for each timepoint. A rolling average consisting of 5 timepoints (250 ms) was then calculated for each animal and pooled together to create the average instantaneous displacement of the vulval muscles around the time of injection (**Figure S1J**). To determine overall displacement caused by injections, the cumulative sum of the average instantaneous displacements was plotted during the 3-second injection window (**Figure S1J** inset).

##### Vulval muscle displacement via prodding

To calculate the displacement caused by the prodding probe, the vulval muscle centroid position was also used. However, since animals were moved by the motorized stage into the stationary prodding needle, the displacement distance observed during prodding was subtracted from the expected displacement had the needle been absent, resulting in the actual displacement upon the vulval muscles. Stage movements were confirmed by expression of mCherry in non-vulval muscle cells from the *unc-103e* promoter (data not shown).

#### Embryo volume estimation

To estimate the volume of an embryo, brightfield images of injection-induced egg-laying events were collected, and embryos (n = 7) were measured for the lengths of the major (length) and minor (diameter) axes. Measurements were averaged and the diameter was calculated to be 27 µm while average length was 51 µm. Calculated embryo volume was ~30 pL when approximated as a cylinder and ~20 pL when approximated as an ellipsoid. Both measurements were averaged to estimate an approximate average volume per embryo of 25 pL.

#### Pharmacological assays

##### Nemadipine

Nemadipine-B (Toronto Research Chemicals Inc.) was dissolved in DMSO to 10 mM and added to melted NGM at 25 µM final concentration prior to media solidification. Control plates containing an equivalent volume of DMSO were made on the same day (0.1% DMSO (v/v) final concentration). Plates were seeded one day prior to the experiment with a 10-fold concentrated OP50 bacteria cultured in B broth. Worms were placed on nemadipine or DMSO plates for at least 1 h prior to microinjections but no longer than 2 h to prevent excess egg accumulation.

##### Floxuridine

Animals were sterilized by exposure to floxuridine as described.^3,16^ In this assay, L4 animals were staged onto NGM plates seeded with OP50, 100 µL of floxuridine (10 mg/mL) was added directly on top of the staged worms and OP50, and animals were incubated at 20 °C for 24 h. Animals were then visually inspected for lack of embryo accumulation prior to selection for microinjection.

### QUANTIFICATION AND STATISTICAL ANALYSIS

#### Data analyses

Injection-induced Ca^2+^ transients were analyzed for the following characteristics: time to peak, amplitude of peak, time to decay from peak, and whether egg laying occurred. We defined ‘Time to peak’ as the time elapsed between the start of injection onset and when the Ca^2+^ trace had reached its peak GCaMP5/mCherry (ΔR/R) value. ‘Peak amplitude’ was defined as the peak value (ΔR/R). ‘Decay time’ was defined as the time when cytosolic Ca^2+^ levels declined to 37% of their peak value (~1/e). Individual Ca^2+^ transient peaks were detected using a custom MATLAB script, as described^64^ and then confirmed via visual inspection of the GCaMP5/mCherry traces from fluorescence recordings. Although multiple injections per animal were often performed, quantitative analyses were performed only on the responses elicited by the first injection. Data were omitted from analyses only for cases in which liquid delivery failed (due to a clogged needle or spillage outside of animal).

#### Statistical analyses

Results were pooled over several days from at least 20 animals (unless otherwise noted). No explicit power analysis was performed prior to the study, but these numbers are in line with prior behavior experiments.^6,10,67^ Statistical analyses were performed using Prism 8 (GraphPad). Isogenic control animals were always tested alongside mutants to account for possible batch effects. Animals were excluded from analyses when liquid delivery failed to spread within the worm (visually determined from brightfield recordings). To determine if there were any statistical differences between the Ca^2+^ responses elicited by injections between wild-type and mutant worms, a Student’s T-Test was performed. In cases in which multiple genotypes or conditions were compared, the data were analyzed by one-way ANOVA. The distributions of responses were tested for normality, and in cases in which data were found to be non-normal, a nonparametric equivalence test was performed (e.g., Mann-Whitney or Kruskal-Wallis). All tests were corrected for multiple comparisons (Bonferroni for ANOVA, Dunn for Kruskal-Wallis). Details of which test was performed for specific strains can be found in figure legends.

##### Standard least squares regression for vulval muscle Ca^2+^ transient amplitude

To determine which factor, as a consequence of injection, best predicted the vulval muscle Ca^2+^ response, a standard least squares regression analysis was performed using JMP Pro 15 (SAS Institute Inc.) with ‘time to peak’, ‘decay time’, ‘number of eggs in uterus’, and ‘flow rate’ as parameters. From this analysis, only ‘number of eggs in uterus’ significantly correlated with vulval muscle Ca^2+^ transient amplitude (p = 0.0095) while other parameters showed a nonsignificant effect. This p-value is represented in **Figure S1B** of which the comparison between ‘number of eggs in uterus’ and ‘vulval muscle Ca^2+^ transient amplitude’ is the only one shown.

##### Nominal logistic fit for egg laying

A similar analysis was performed using JMP to determine which factor best predicted egg release following injection. The parameters used were ‘vulval muscle Ca^2+^ transient amplitude’, ‘flow rate’, ‘time to peak’, ‘decay time’, and ‘number of eggs in uterus’. Only ‘vulval muscle Ca^2+^ transient amplitude was a significant parameter (p < 0.0001) while other parameters provided a nonsignificant effect.

### KEY RESOURCES TABLE

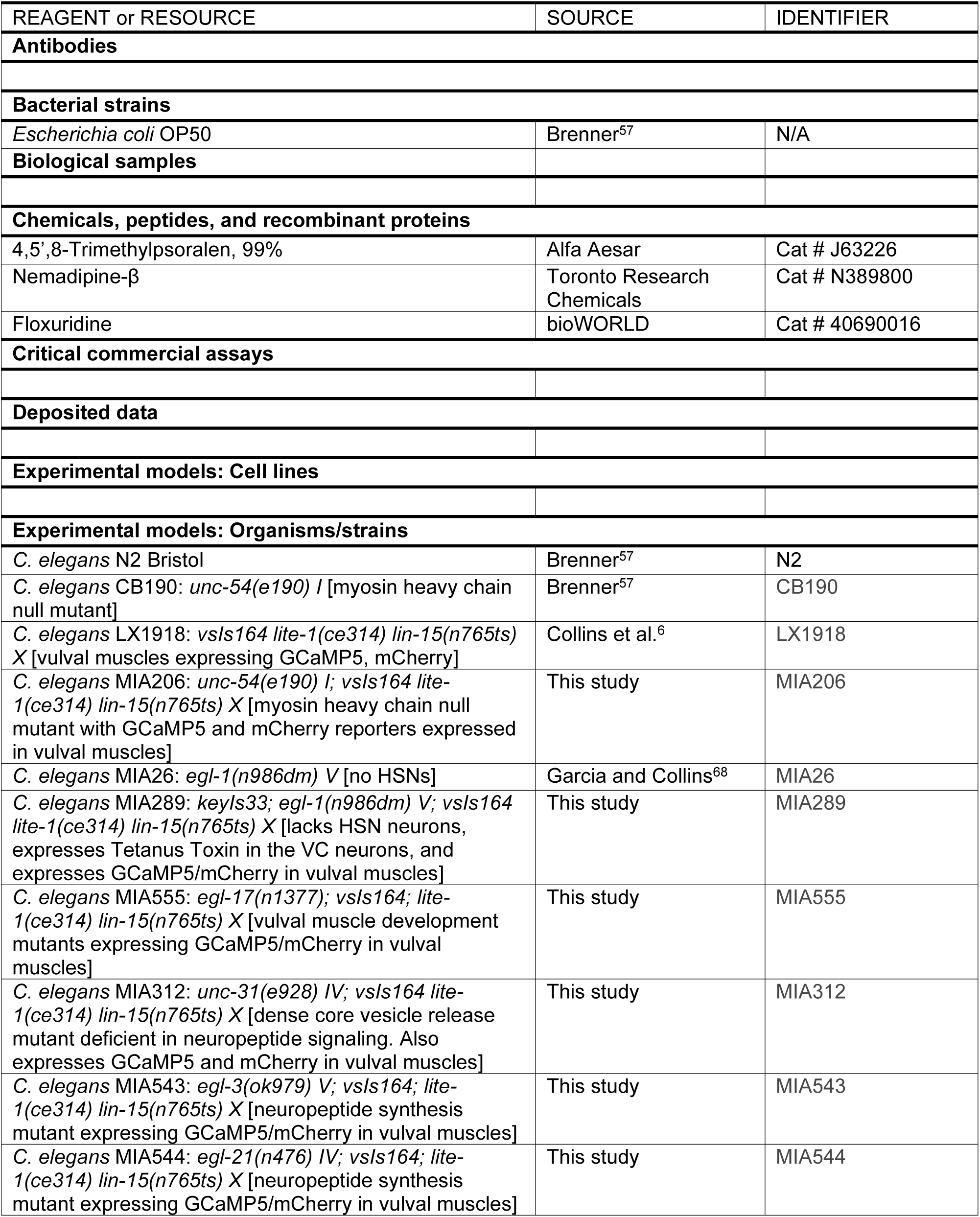
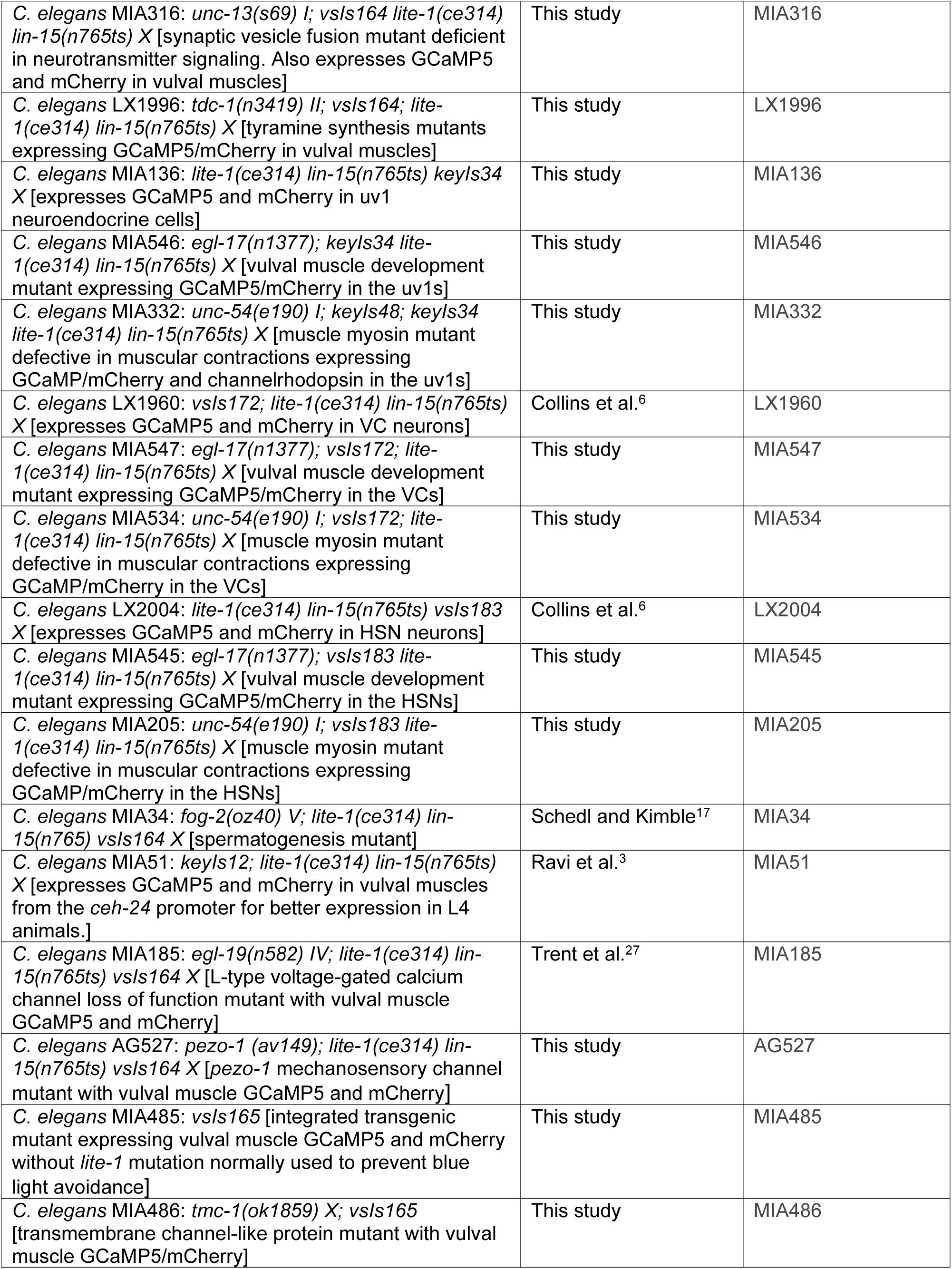
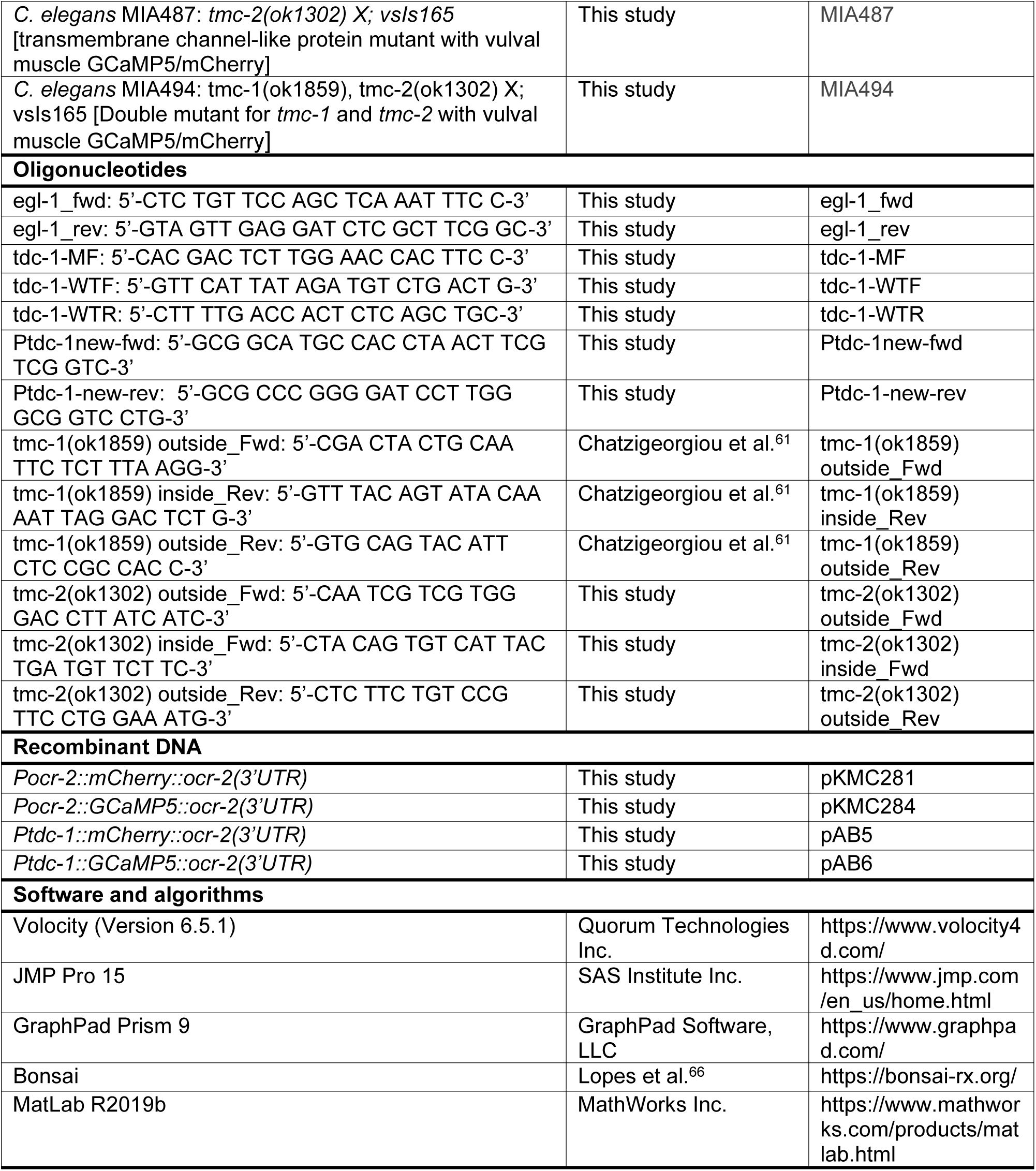

## Legends for supplemental videos

**Video S1. Microinjection with bromophenol blue dye into vulval muscle GCaMP reporter strain. Related to Figure 1 and Figure S2.**

Recording of vulval muscle GCaMP5 (green) and bromophenol blue (magenta) fluorescence during microinjection overlayed over a brightfield recording of *C. elegans.* Wild-type animals display an increase in vulval muscle (vm) Ca^2+^ activity that correlates with the increase in injection volume (represented as traces within the video). Egg release is coincident with peak vulval muscle GCaMP5 fluorescence.

**Video S2. Microinjection into vulval muscle GCaMP reporter strain exposed to 25 µM nemadipine or 0.1% DMSO. Related to Figure 1.**

GCaMP5 to mCherry fluorescence ratio in vulval muscles of a DMSO-treated control animal (top) and a nemadipine-treated animal (bottom) overlaid on top of respective brightfield recording during injection. Blue indicates low Ca^2+^, and red indicates high Ca^2+^. Red traces denote vulval muscle Ca^2+^ responses and white horizontal bar denotes timing of the 3 s injection pulse. Inserts on right side of movie show a zoomed in recording of vulval muscle Ca^2+^ response for control and Nemadipine-treated animals to emphasize localization of Ca^2+^ activity. White arrowhead shows localization of residual Ca^2+^ activity from vm1 muscles in nemadipine-treated animals.

**Video S3. Microinjection into uv1 GCaMP reporter strain. Related to Figure 2.**

GCaMP5 to mCherry fluorescence ratio in uv1 cells of wild-type animals overlaid on top of brightfield recording during injection. Blue indicates low Ca^2+^, and red indicates high Ca^2+^. Red trace denotes uv1 Ca^2+^ response, triangle denotes egg release, and white horizontal bar denotes timing of the 3 s injection pulse.

**Video S4. Microinjection into VC GCaMP reporter strain. Related to Figure 2.**

GCaMP5 to mCherry fluorescence ratio in VC neurons of wild-type animals overlaid on top of brightfield recording during injection. Blue indicates low Ca^2+^, and red indicates high Ca^2+^. Red trace denotes VC Ca^2+^ response, triangle denotes egg release, and white horizontal bar denotes timing of the 3 s injection pulse.

**Video S5. Microinjection in HSN GCaMP reporter strain. Related to Figure 2.**

GCaMP5 to mCherry fluorescence ratio in HSN neurons of wild-type animals overlaid on top of brightfield recording during injection. Blue indicates low Ca^2+^, and red indicates high Ca^2+^. Red trace denotes HSN Ca^2+^ response, triangle denotes egg release, and white horizontal bar denotes timing of the 3 s injection pulse.

**Video S6. Prodding of vulval muscles. Related to Figure 4.**

Recording of vulval muscle mCherry (magenta) and vulval muscle GCaMP5 (green) fluorescence during mechanical prodding of the posterior but not anterior vulval muscles. Cyan outline border of the prodding needle. White horizontal bar denotes timing of 1 s prodding stimulus.

